# Gene Therapy Mediates Therapeutic Improvement in Cardiac Hypertrophy and Survival in a Murine Model of *MYBPC3*-Associated Cardiomyopathy

**DOI:** 10.1101/2024.02.19.581102

**Authors:** Amara Greer-Short, Anna Greenwood, Elena C. Leon, Tawny Neal Qureshi, Konor von Kraut, Justin Wong, Christopher A. Reid, Ze Cheng, Emilee Easter, Jin Yang, Jaclyn Ho, Stephanie Steltzer, Ana Budan, Marie Cho, Charles Feathers, Tae Won Chung, Neshel Rodriguez, Samantha Jones, Chris Alleyne-Levy, Jun Liu, Frank Jing, William S. Prince, JianMin Lin, Kathryn N. Ivey, Whittemore G. Tingley, Timothy Hoey, Laura M. Lombardi

## Abstract

**Background:** Hypertrophic cardiomyopathy (HCM) affects an estimated 600,000 people in the U.S. and is the leading cause of sudden cardiac arrest in those under 18. Loss-of-function mutations in *Myosin Binding Protein C3*, *MYBPC3*, are the most common genetic cause of HCM. The majority of *MYBPC3* mutations causative for HCM result in truncations. The sarcomeric pathophysiology of the majority of HCM patients with *MYBPC3* mutations appears to be due to haploinsufficiency, as the total amount of MYBPC3 protein incorporated into sarcomeres falls significantly below normal.

**Methods:** A clear path for the treatment of haploinsufficiency is the restoration of the insufficient gene product; in this case wild-type MYBPC3. To achieve this, we engineered an AAV vector (TN-201) with superior properties for mediating cardiomyocyte-selective expression of MYBPC3 after systemic delivery.

**Results:** We have demonstrated for the first time with AAV gene therapy the ability of both a mouse surrogate and TN-201, which encodes human MYBPC3 to reverse cardiac hypertrophy and systolic dysfunction and to improve diastolic dysfunction and survival in a symptomatic MYBPC3-deficient murine model of disease. Dose-ranging efficacy studies exhibited restoration of wild-type MYBPC3 protein levels and saturation of cardiac improvement at the clinically relevant dose of 3E13 vg/kg, outperforming a previously published construct. Further, we have established stable cardiac benefit for greater than one year post-injection, as well as reversal of cardiac dysfunction even in late-stage models of disease.

**Conclusions:** Our data suggest that by restoring MYBPC3 to the sarcomere, TN-201 has the potential to slow and even reverse the course of the disease in patients with *MYBPC3*-associated HCM.

## INTRODUCTION

Mutations in *MYBPC3*, the gene encoding cardiac myosin-binding protein C (MYBPC3 or cMyBP-C), are the most common genetic cause of HCM, an autosomal dominant condition.^1^ HCM increases the risks of sudden cardiac death (SCD) and is a leading cause of SCD in children and young adults.^2–5^ In primary literature, the reported prevalence of *MYBPC3* mutations in patients with diagnosed HCM ranges from 17–26%.^1,6–10^ *MYBPC3*-associated HCM is a progressive disease characterized by left ventricular (LV) hypertrophy, small LV volume, ventricular diastolic dysfunction, cardiac arrhythmias, and imbalance between myocardial oxygen supply and demand. Disease burden may include dyspnea, exercise intolerance, atypical angina, syncope, ventricular arrhythmias, SCD, and heart failure. The Sarcomeric Human Cardiomyopathy Registry (SHaRe) identified risk factors for HCM severity based on genotype and lifetime burden of disease. Primary risk factors included an identified sarcomere mutation (including *MYBPC3*) and early onset of disease (before age 40 years).^1^

Cardiac MYBPC3 protein has structural and functional roles in sarcomere biology, acting as a brake that limits actin-myosin crossbridge interactions during cardiac contraction and relaxation.^11–14^ *MYBPC3* truncating mutations reduce MYBPC3 protein content, thereby enhancing maximal thin filament sliding velocity within the thick filament C zone. The pathophysiology of *MYBPC3*-associated HCM is clearly attributable to MYBPC3 protein deficiency. Myectomy samples from patients who have HCM with a *MYBPC3* mutation had 40% lower MYBPC3 protein levels, with no detectable expression of truncated MYBPC3 from the mutant *MYBPC3* gene, compared to normal control samples.^15–19^ The lack of detectable truncated MYBPC3 suggests that either no protein was synthesized from the mutant *MYBPC3* allele or, alternatively, that the truncated protein was susceptible to rapid degradation. Truncating variants account for 91% of pathogenic *MYBPC3* genes and cause similar clinical severity and outcomes regardless of the location of the truncation within the protein product, consistent with locus-independent loss of function.^20^ These data support a rationale for developing TN-201, a *MYBPC3* gene replacement therapy, which is designed to correct both the pathogenic reduction in MYBPC3 in heterozygotes and the total loss of protein in homozygotes and compound heterozygotes,^21–23^ which can exhibit severe dilatation and typically experience heart failure and death within the first year of life.^24^

Gene therapy is potentially curative and increasingly feasible for a myriad of genetic and chronic diseases. Although AAV currently represents the most tractable viral vehicle due to superior safety and manufacturability, the size constraints of the AAV genome (4.7 kb) frequently limit which therapeutic cargoes can be delivered and the extent to which they are expressed.^25–27^ Robust transgene expression is often dependent on the inclusion of two intact flanking ITR sequences, a promoter, intron, WPRE, and polyadenylation signal. Thus, standard *cis*-regulatory sequences used in AAV transgene expression cassettes amount to ∼1.2 kb without the promoter. This limits the ability to efficiently package large genes which require tissue-specific expression in AAV, including our primary therapeutic target *MYBPC3* cDNA (3.825 kb).

The purpose of this study was to engineer an optimal AAV genomic cassette for MYBPC3 expression in cardiomyocytes and test in an *in vivo* disease model dependent on MYBPC3 loss. We demonstrated for the first time durable improvement in cardiac phenotypes, including reduction of cardiac hypertrophy, improvement in systolic and diastolic function, and extended survival in a murine model of *MYBPC3*-associated cardiomyopathy.

## METHODS

Full Methods are described in the Supplemental Material.

### Study design

The number of biological and technical replicates (N) per experiment is noted in each figure legend. Mice were randomized before being assigned to either vehicle or treatment groups. An individual blinded to the treatment performed and analyzed the echocardiography data.

### iPSC-CM generation

*MYBPC3^−/−^* iPSCs were generated by a CRISPR-Cas9 paired gRNA approach to remove exons one and two from WTC iPSCs. Differentiations into iPSC-CMs were conducted as previously described.^28^ Briefly, differentiation of iPSCs was induced through the Wnt modulation method with 7 μM glycogen synthase kinase 3 b (GSK3b) inhibitor CHIR99021 (Selleckchem) and 5 μM inhibitor of WNT production 2 (IWP2) (Sigma). Basal Media was composed of RPMI media supplemented with B27 (Thermo Fisher Scientific). Eighteen days after adding CHIR, iPSC-CMs were frozen in 90% fetal bovine serum with 10% dimethyl sulfoxide (DMSO).

### AAV production

AAV production was carried out as previously described.^29^ Briefly, HEK293T cells were seeded in a Corning HyperFlask (New York, NY) and triple transfected using a 2:1 PEI:DNA ratio (PEI Max) with a helper plasmid containing adenoviral elements (pHelper), a plasmid containing the Rep2 and Cap genes from the respective AAV serotype, and finally an ITR containing plasmid to be packaged. Three days following transfection cells were harvested and lysed. Virus was purified using iodixanol ultracentrifugation and cleaned and concentrated in Hank’s Balanced Salt Solution (HBSS) + 0.001% Pluronic using a 100-kDa centrifuge column (Amicon, Darmstadt, Germany). AAV was titered using a PicoGreen assay.

## RESULTS

### Engineering a Leaner AAV Genomic Cassette without Loss of Strength or Specificity of Cardiac Expression

The size constraints of the AAV genome are highly restrictive for optimal expression of large therapeutic genes which require tissue-specific expression, such as *MYBPC3* (ORF = 3.825 kb). After determining that only minor protein expression differences occurred upon deletion of the WPRE element (540 bp) and minimization of sequences required for polyadenylation (Figure S1), we focused our attention on minimizing a cardiac promoter without loss of strength or specificity for *MYBPC3* gene replacement therapy.

We first established successful transgene expression and protein localization in *MYBPC3*^−/−^ induced pluripotent stem cell-derived cardiomyocytes (iPSC-CMs) (Figure 1A). We then tested multiple versions of the minimized cassette (compared to a 5.6 kb standard cassette) with different cardiac promoters, called pCard, driving expression of human *MYBPC3* (Figure 1B). The constructs were packaged into AAV, used to infect human *MYBPC3^−/−^*iPSC-CMs and their resultant protein expression ascertained by immunoblotting. Surprisingly, a smaller version of the pCard0 promoter, pCard1, drove the greatest expression (Figure 1C and 1D; from the human cardiac human cardiac troponin T (*TNNT2*) promoter; sequence in patent US-20210252165-A1). This is likely due to increased packaging of the AAV genome, since the pCard0 construct was 4.9 kb, exceeding the size of the wild-type AAV genome, 4.7 kb. Consistent with the pCard1 4.7 kb version driving increased expression through improved packaging, no difference in protein expression was observable when *MYBPC3^−/−^* iPSC-CMs were transiently transfected with the naked plasmids (Figure S2A).

**Figure 1.**
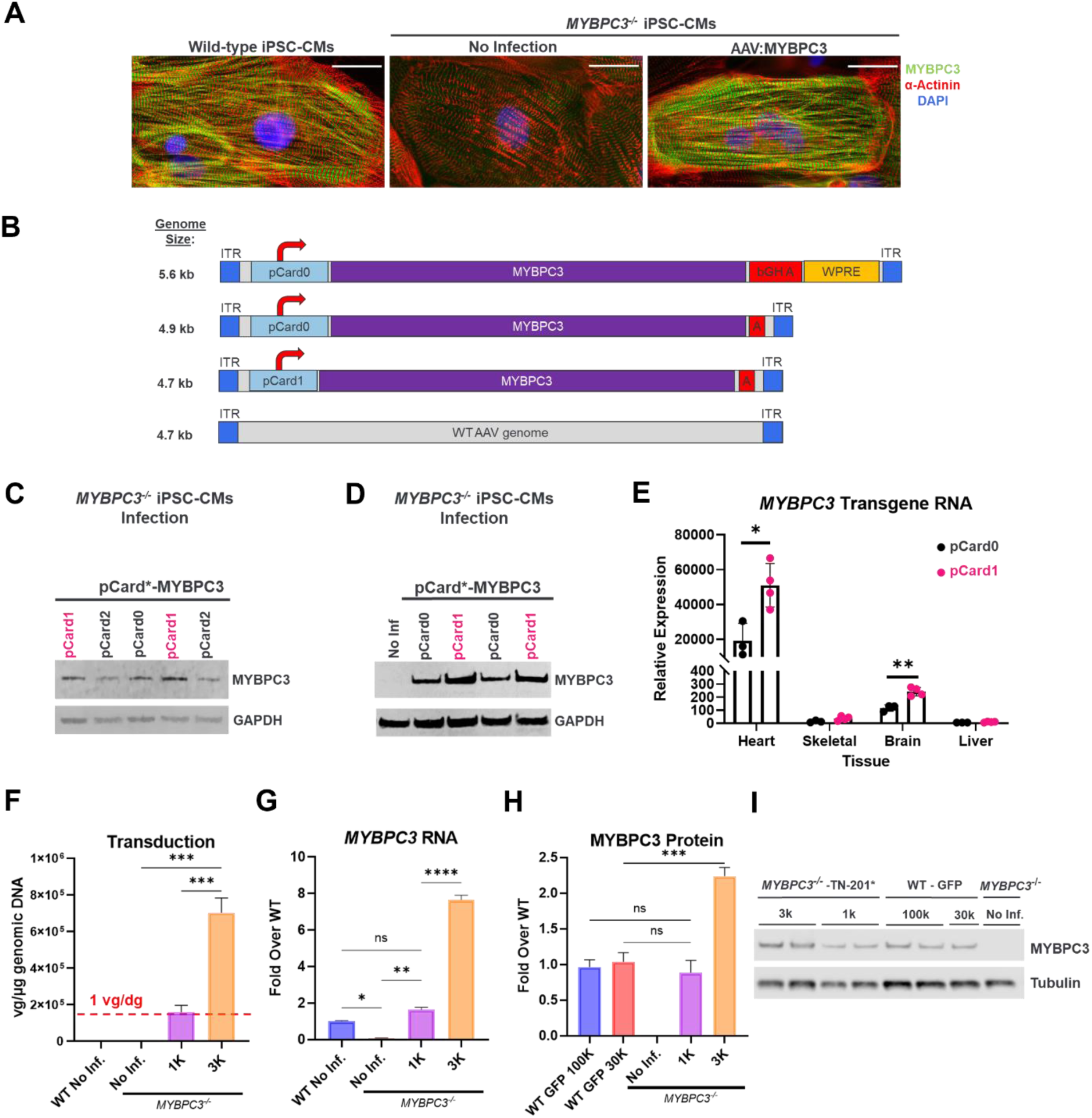
AAV Genomic Cassette and Promoter Engineering to Drive Potent and Specific Expression. (**A**) In order to validate the assay system, we confirmed transgene expression resulted in proper localization in human *MYBPC3^−/−^* iPSC-CMs transduced with AAV:MYBPC3. Immunofluorescence analysis was performed seven days post-infection (scale bars, 25 µm). (**B**) Cassette schematic indicating the genomic size of a standard cassette and the alterations tested. (**C**) Human *MYBPC3^−/−^*iPSC-CMs were transduced with AAV6-packaged constructs encoding human *MYBPC3* driven by various promoter versions of human cardiac troponin T *(TNNT2)* (pCard0, pCard1 and pCard2) and harvested 5 days post-infection (n = 1-2/condition). (**D**) A head-to-head comparison of the independently packaged versions of the pCard0 and pCard1 constructs. *MYBPC3^−/−^* iPSC-CMs were harvested five days post-infection with CR9-01-packaged constructs (n = 2/condition). (**E**) To determine if the optimized promoter also increased expression in vivo and whether selectivity was maintained, adult mice were retro-orbitally injected with 4E13 vg/kg AAV9 encoding the pCard0 and pCard1 constructs. Heart, skeletal muscle (tibialis anterior), liver and whole brain samples were harvested two weeks post-injection. (n = 3-4/condition; P-value per Student’s t-test, * p < 0.5, ** p <0.01). Matched transduction (**F**), RNA (**G**) and protein (**H**) analysis in *MYBPC3*^−/−^ iPSC-CMs demonstrated near-WT levels of protein expression at 1 vg/dg, with (**I**) representative immunoblot in *MYBPC3*^−/−^ iPSC-CMs compared to parental wild-type control iPSC-CMs one week post-infection (n = 2-3/condition assayed; one-way ANOVA with Tukey’s multiple comparisons test, * p < 0.5, ** p <0.01, *** p<0.001).. Data are shown as means ± SEM.

Critical to the utility of this shorter and stronger promoter is retention of cardiac selectivity. We therefore packaged the pCard0 and pCard1 versions into AAV9 for assessment of strength and selectivity *in vivo*. Adult mice were retro-orbitally injected with 4E13 vg/kg and tissue samples from heart, skeletal muscle (tibialis anterior), liver and whole brain were harvested two weeks post-injection. RNA was extracted from all tissues and analyzed by qPCR with primers specific to human *MYBPC3*. Both viral preps resulted in robust RNA expression in the heart, with ∼200X selectivity of expression in heart over brain and >1000X selectivity over skeletal muscle and liver (Figure 1E). Thus, the boost in expression of the pCard1 construct was confirmed *in vivo* (>2x improvement in cardiac RNA expression) and, critically, this decrease in size did not affect the selectivity of expression *in vivo*. Thus, AAV9 packaging pCard1-MYBPC3 was selected for further development as a therapeutic candidate and henceforth designated TN-201.

To evaluate the expression strength of the TN-201 genomic cassette in human cardiomyocytes, homozygous mutant *MYBPC3*^−/−^ iPSC-CMs were generated and the dose-responsiveness of TN-201 genomic cassette-mediated transgene expression was compared with MYBPC3 levels in WT isogenic control iPSC-CMs. To achieve the greatest levels of iPSC-CM infection for this transduction assessment, the TN-201 AAV genomic cassette was packaged into the AAV9 serotype variant CR9-01, which exhibits higher infectivity than AAV9 of cultured iPSC-CMs. This enabled efficient utilization of the virus, while maintaining the integrity of TN-201 AAV genomic cassette assessment. The CR9-01:hMYBPC3 treatment is henceforth denoted as TN-201* to distinguish it from TN-201. *MYBPC3* transgene RNA and MYBPC3 protein levels in *MYBPC3*^−/−^ iPSC-CMs were compared with endogenous levels of *MYBPC3* RNA and MYBPC3 protein in WT iPSC-CMs one week post-infection.

To precisely determine how these transgene expression evaluations mapped to transduction levels, matched vector genome (Figure 1F), RNA (Figure 1G), and protein (Figure 2H and 2I) analyses were performed one week post-infection. Absolute quantification of nuclear vector genomes indicated that the lowest dose administered (1K TN-201*) resulted in ∼1.6E5 vg/µg DNA, which is equivalent to ∼1 vector genome per diploid genome (vg/2n). Matched assessment of *MYBPC3* RNA and MYBPC3 protein at this dose indicated ∼1.6X the endogenous level of *MYBPC3* RNA and ∼0.9X the endogenous level of MYBPC3 protein. Consistent with vector genome analysis, flow cytometry one week post-infection with CR9-01:pCard1-GFP, a green fluorescent protein reporter, demonstrated that at the lowest dose used (1K) ∼60% of iPSC-CMs were transduced (Figure S2B), indicating an infectious multiplicity of infection equivalent to one based on the Poisson distribution.^30^ Thus, transduction equivalent to approximately one vector genome per diploid genome drove near WT levels of *MYBPC3* RNA and MYBPC3 protein one week post-infection. This strength of expression in the hiPSC-CM context propelled us to move forward in attempting to treat an *in vivo* model of *MYBPC3*-associated cardiomyopathy.

**Figure 2.**
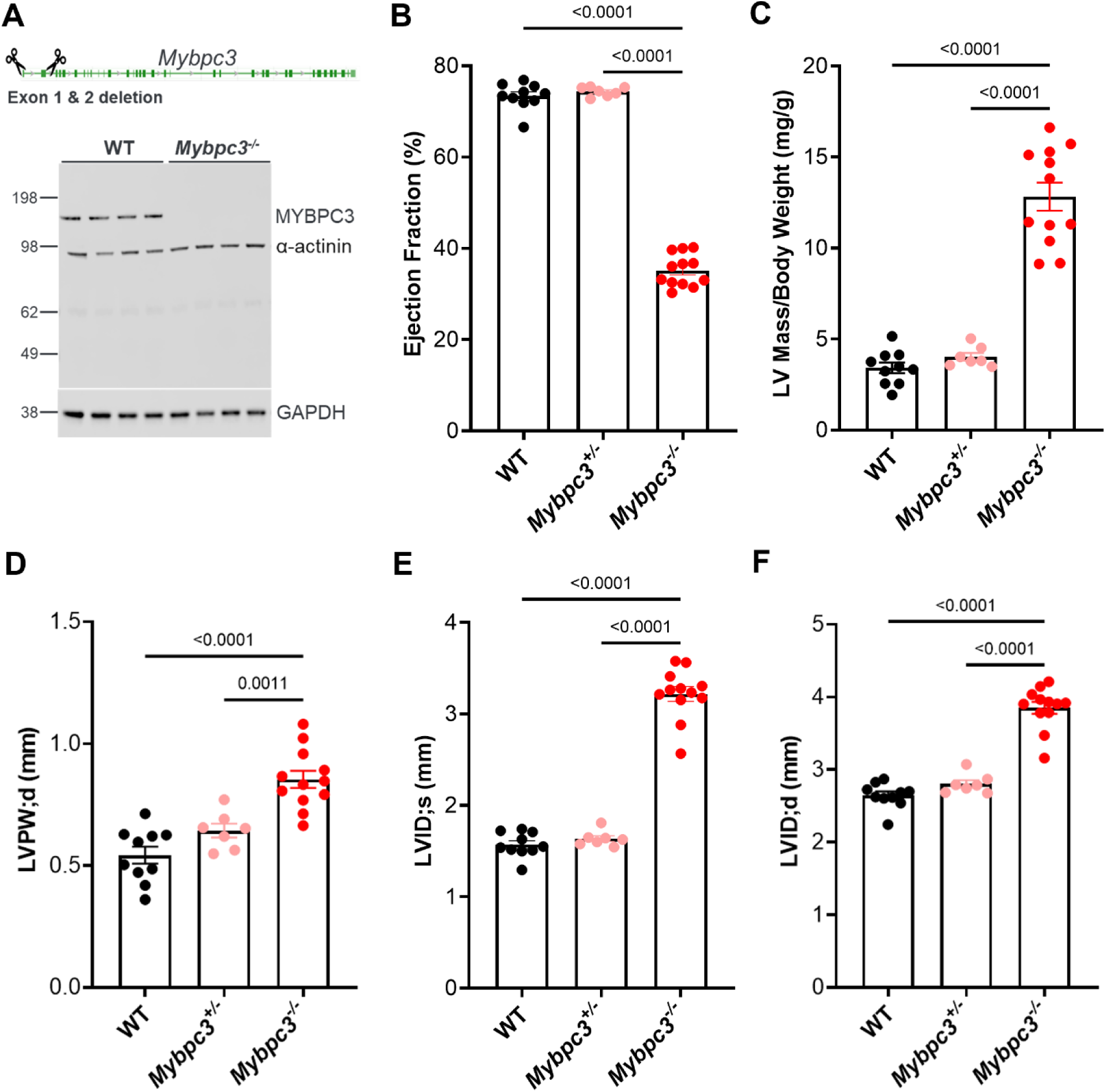
Murine *Mybpc3^−/−^* Model for Treatment of HCM. (**A**) A null mutant allele of *Mybpc3* was generated by excision of exons one and two using CRISPR-Cas9 and confirmed by demonstrating MYBPC3 depletion in an immunoblot from cardiac tissue (n = 4/condition). Echocardiography analysis of homozygous mutants, heterozygous mutants and WT littermates demonstrated a significant decrease in cardiac systolic function in homozygous animals based on (**B**) ejection fraction (%) at two weeks of age. (**C**) Homozygous mice exhibited marked LV hypertrophy at two weeks of age as evidenced by their LV mass normalized by body weight and (**D**) increased left ventricular posterior wall thickness during diastole (LVPW;d). Left ventricular internal diameters during (**E**) systole (LVID;s) and (**F**) diastole (LVID;d) were also significantly increased in homozygotes at two weeks of age. WT (n = 10), *Mybpc3^+/−^* (n = 7) and *Mybpc3*^−/−^ (n = 12). Heterozygous mice were not significantly different from WT in any parameter. P-value per one-way ANOVA with Tukey’s multiple comparisons test. Data are shown as means ± SEM.

### Murine *Mybpc3^−/−^* Model for Treatment of HCM

*Mybpc3^−/−^* mice were established on the C57BL/6 background as a model for gene replacement of *MYBPC3* in severely affected HCM patients by a CRISPR-Cas9 paired gRNA deletion of exons one and two (Figure 2A; please refer to Supplemental Methods). Heterozygous and homozygous animals were compared with WT controls for a number of cardiovascular function and morphology parameters. Heterozygous *Mybpc3^+/−^* mice showed no cardiovascular phenotype and were therefore not considered to be a useful model for efficacy studies. Homozygous mice exhibited severe deficits in cardiac function (Figure 2B) and pronounced cardiac hypertrophy (Figure 2C through 2F) as early as two weeks of age (Table S1). With body weight normalized LV mass 3-4 times that of WT siblings, increased posterior wall thickness or LVPW;d (1.6x WT siblings), and severe cardiac dysfunction (ejection fraction (EF): 35% to 40%), this model closely mimics late-stage HCM, as well as in infants with compound heterozygous or homozygous mutations in the *MYBPC3* gene. LV hypertrophy is the defining feature of all patient classes with wall thickness as a key prognostic marker for HCM and a major risk factor for cardiovascular mortality and SCD.^31^ LV dilatation is more common in pediatric compound heterozygotes or homozygotes.^24,32–35^ Due to the severe phenotype of the *Mybpc3^−/−^* mice, and the lack of any MYBPC3 protein, this is a challenging model to demonstrate efficacy.

### AAV9:mMybpc3 and TN-201 Improved Hypertrophy, Cardiac Dysfunction and Premature Lethality of *Mybpc3^−/−^* Mice

In order to test whether AAV treatment can improve the cardiac phenotype of *Mybpc3^−/−^* mice with established disease, and not merely prevent disease progression when injected before the onset of cardiac symptoms,^36^ AAV9:mMybpc3, the mouse surrogate of TN-201 (also known as mTN-201), and TN-201 were administered to symptomatic *Mybpc3^−/−^* mice. To our knowledge, this experiment represents the first *in vivo* assessment of the human MYBPC3 activity in a mouse model. The mouse ortholog was utilized for the appropriate species-matched comparison, since the human ortholog is only 88% identical at the amino acid level. Vehicle, AAV9:mMybpc3, or TN-201 were administered systemically (1E14 vg/kg) via retro-orbital intravenous (IV) injection to symptomatic 2-week-old *Mybpc3^−/−^* mice (9-12/group), when the EF averaged 35.2 ± 1.0% (versus 73.4 ± 0.9% in WT mice) and normalized LV mass was 4-fold above WT. An additional group of 12 WT littermates were administered vehicle as a control. Echocardiography was performed to assess cardiac function until Week 54. Animal survival was followed up until animals were 20 months of age, at which time the remaining animals were euthanized.

Both treatments significantly improved LV hypertrophy, cardiac function, and lifespan relative to vehicle-treated *Mybpc3^−/−^* mice. As might be anticipated based on sequence divergence between the mouse and human genes, the species-matched treatment, AAV9:mMybpc3, produced larger improvements than treatment with the human ortholog, TN-201. Both AAV9:mMybpc3 and TN-201 significantly decreased LV hypertrophy (Figure 3A), with mean improvements of 5.1 ± 0.7 mg/g and 4.2 ± 0.8 mg/g for LV mass respectively, compared to vehicle-treated *Mybpc3^−/−^* mice at 31 weeks following viral delivery (Figure 3B and Table S2). Treatment with AAV9:mMybpc3 increased EF by up to 26% above baseline over 6 weeks and then had a stable effect on EF up until the animals were 13 months of age, which was the age of the last echocardiography measurements. Animals treated with TN-201 had a 5% increase in EF over 10 weeks from baseline and then showed a slow decline in EF as they aged (Figure 3C). Both AAV9:mMybpc3 and TN-201 resulted in superior cardiac function compared to vehicle-treated *Mybpc3^−/−^* mice, with mean EF improvements of 30 ± 1.7% and 18 ± 1.9%, respectively, at 31 weeks following viral delivery (Figure 3D and S3).

**Figure 3.**
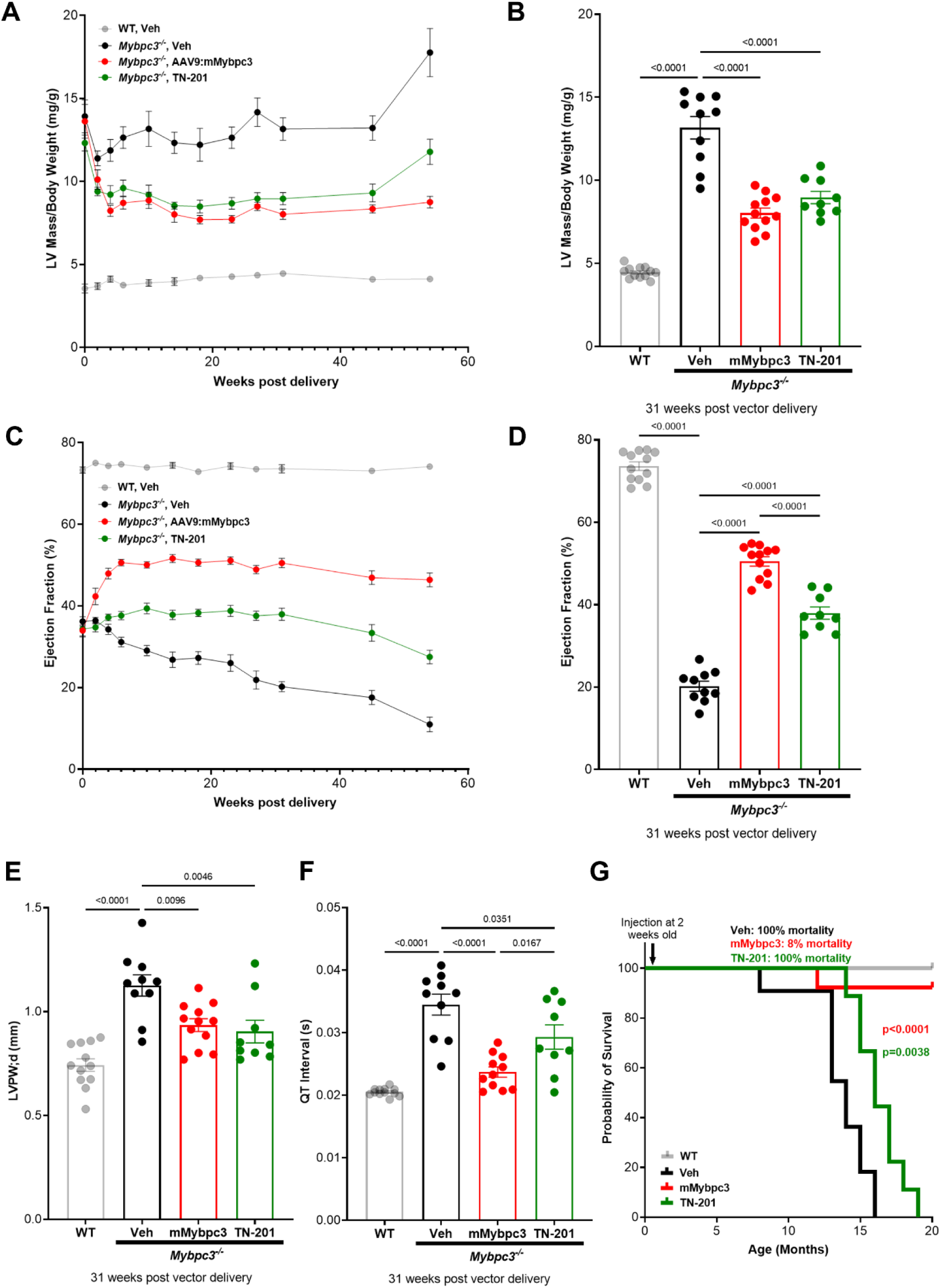
AAV9:mMybpc3 and TN-201 Improved Hypertrophy, Cardiac Dysfunction and Premature Lethality of *Mybpc3*^−/−^ Mice. (**A**) LV mass normalized to body weight progression over time and (**B**) at 31 weeks post-delivery. (**C**) EF progression and (**D**) EF at 31 weeks post-delivery. **(E)** Left ventricular posterior wall thickness at the end of diastole (LVPW;d) and (**F**) QT interval at 31 weeks post-delivery. (**G**) Kaplan-Meier survival curve with animals followed out until 20 months of age. Median survival for *Mybpc3^−/−^* vehicle animals was 14 months, and all animals died by 16 months of age. For TN-201-treated animals, median survival was 16 months (i.e. 2 months lifespan extension from vehicle), and all animals died by 19 months of age. AAV9:mMybpc3 treatment extended lifespan, with only one animal euthanized due to skin lesions not related to heart failure by 20 months. Data are shown as means ± SEM. WT mice were significantly different from all groups for all parameters, with the exception of AAV9:mMybpc3-treated animals for QT interval. P-value per one-way ANOVA with Tukey’s multiple comparisons test.

Notably, AAV9:mMybpc3 and TN-201 significantly improved posterior wall thickness by 0.19 ± 0.06 mm and 0.22 ± 0.07 mm, respectively, relative to vehicle (Figure 3E) at 31 weeks post-delivery and was again indicative that both treatments improve LV hypertrophy. Similar to patients with *MYBPC3*-associated HCM^37^ and consistent with established mouse models,^38,39^ *Mybpc3^−/−^* mice exhibit QT interval prolongation. Critically, both treatments significantly improved QT prolongation compared to vehicle-treated *Mybpc3^−/−^* mice (Figure 3F) and lifespan was significantly increased in both treatment groups (Figure 3G). AAV9:mMybpc3 treatment resulted in lifespan extension >6 months, with only one animal death prior to study-wide euthanasia, which was due to factors unrelated to heart failure. Even the TN-201 treatment group had their lifespans extended by approximately two months. Thus, species-matched treatment resulted in profound improvements in LV hypertrophy, EF, QT interval prolongation and survival. Further, despite the sequence divergence between the mouse and human orthologs, TN-201 mediated significant efficacy in multiple parameters assessed in this mouse cardiomyopathy model, providing proof of activity for our human gene therapy in an *in vivo* model.

### AAV9:mMybpc3 Improved Cardiac Function at a Dose as Low as 1E13 vg/kg

Given the proof-of-concept study was performed at 1E14 vg/kg, we moved forward with titration of AAV9:mMybpc3 to determine the dose-response relationship for cardiac improvement. We evaluated three single doses of AAV9:mMybpc3 in *Mybpc3*^−/−^ mice. The treatments were given systemically at two weeks of age via retro-orbital injections, at 1E13 vg/kg, 3E13 vg/kg or 1E14 vg/kg. As a control, vehicle was injected to a group of wild type (WT) and a group of *Mybpc3*^−/−^ animals.

AAV9:mMybpc3 was beneficial to the recovery of cardiac function at every dose tested, with dose-dependent improvements in LV mass (Figure 4A), EF (Figure 4B), and QT interval (Figure 4C). The dose of 3E13 vg/kg yielded robust and highly significant improvements with decreased hypertrophy of 5.3 ± 0.8mg/g LV Mass and improved EF of 26 ± 3.0%. A similar trend was observed for improvement of QT prolongation (Figure 4C). The lower dose of 1E13 vg/kg dose produced smaller improvements with decreased hypertrophy of 3.2 ± 1.3 mg/g LV mass and improved cardiac function as assessed by EF of 15 ± 3.7%. Indicative of treatment tolerability in this disease model, there were no differences in body weights among the treatment groups (Figure 4D). Of note, animals treated with 3E13 and 1E14 vg/kg did not exhibit significant differences in improvement of hypertrophy, EF or QT interval prolongation, suggesting a plateau in effect. Thus, the clinically relevant dose of 3E13 vg/kg^40,41^ appeared to achieve near-maximal efficacy for saturation of cardiac benefit.

**Figure 4.**
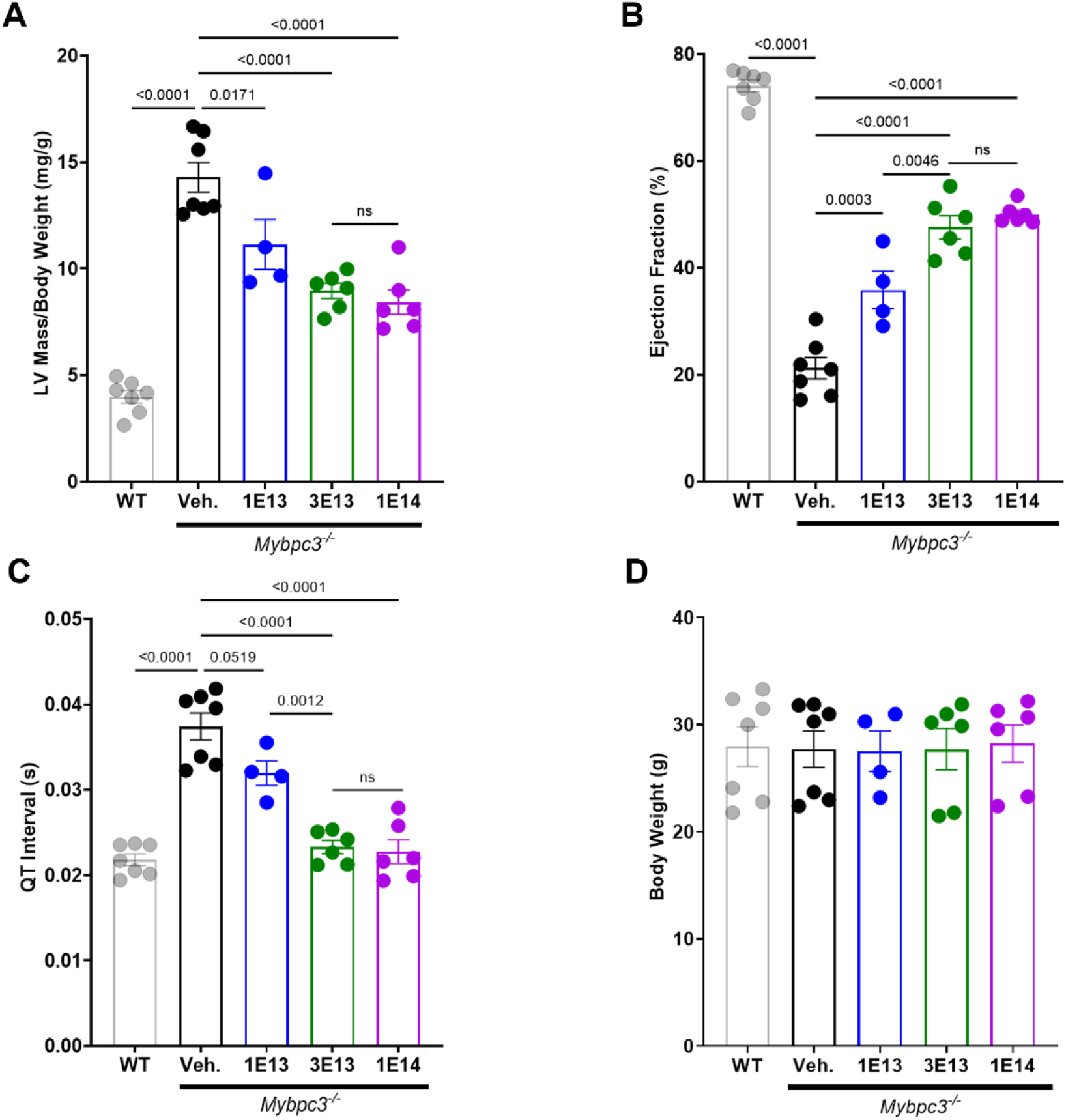
AAV9:mMybpc3 Improves Cardiac Function at a Dose as Low as 1E13 vg/kg. Dose-dependent improvement compared to vehicle-treated *Mybpc3^−/−^* mice in (**A**) LV mass normalized to body weight, (**B**) EF, and (**C**) QT interval at 31 weeks post-delivery. (**D**) No group differences in body weight at 31 weeks post-delivery. Data are shown as means ± SEM. WT mice were significantly different from all groups for all echocardiographic parameters, with the exception of 3E13 vg/kg and 1E14 vg/kgAAV9:mMybpc3-treated animals for QT interval. P-value per one-way ANOVA with Tukey’s multiple comparisons test.

### Cardiac Restoration of Wild-Type MYBPC3 Protein Levels

Given that animals treated with 3E13 and 1E14 vg/kg did not exhibit significant differences in improvement of hypertrophy, EF or QT interval prolongation, we hypothesized that both doses were adequate to restore wild-type levels of MYBPC3 protein to deficient mice. The kinetics and dose-responsiveness of transgene RNA and protein expression in *Mybpc3^−/−^* mice was assessed to match efficacy studies with AAV9:mMybpc3. Doses of 3E13 vg/kg and 1E14 vg/kg were administered systemically using retro-orbital IV injection at two weeks of age. Two weeks post-injection, homozygotes dosed with 3E13 and 1E14 vg/kg exhibited WT levels of cardiac MYBPC3 protein expression, as assessed by immunoblot (Figure 5A and phosphorylation Figure S4) and ELISA (Figure 5B). Similar results were obtained six weeks post-injection, with dose-dependent *Mybpc3* RNA expression observed (Figure 5C).

**Figure 5.**
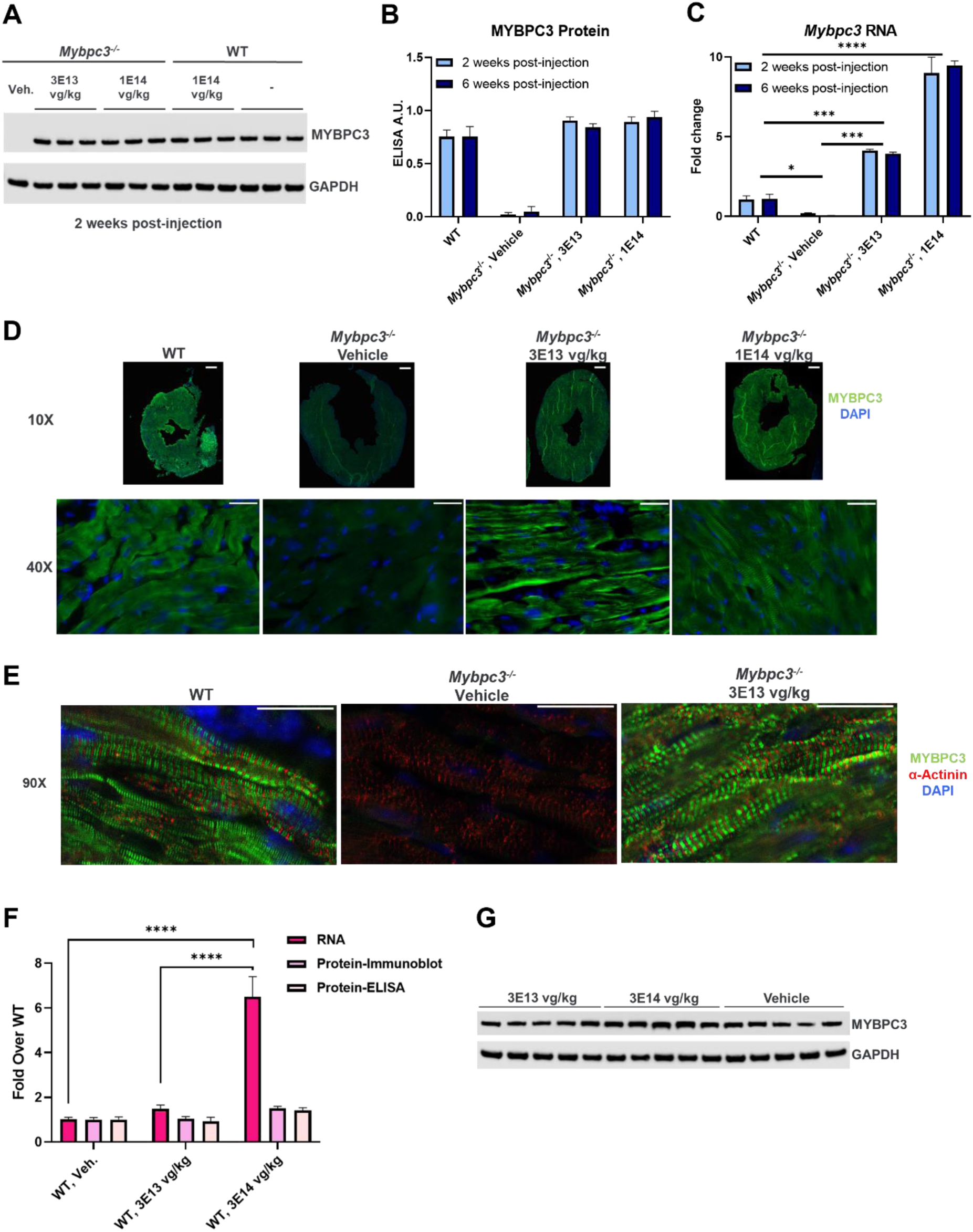
Cardiac Restoration of Wild-Type MYBPC3 Protein Levels. Restoration of WT levels of cardiac MYBPC3 protein upon dosing *Mybpc3*^−/−^ mice at two weeks of age with 3E13 and 1E14 vg/kg based upon (**A**) immunoblot and (**B**) ELISA (n = 2-3/condition). Homozygous mice were dosed retro-orbitally IV with vehicle or the indicated doses (vg/kg) of AAV9:mMybpc3, and cardiac protein analyzed two weeks and six weeks post-injection. (**C**) *Mybpc3* transgene RNA two weeks and six weeks post-injection (n = 2-3/condition). (**D**) Immunohistochemistry for MYBPC3 4 weeks post-injection. Homozygous mice were dosed retro-orbitally IV with vehicle or the indicated doses (vg/kg) of AAV9:mMybpc3 at two weeks of age. WT littermates were harvested, and samples processed simultaneously (n = 2/condition; 10X scale bars: 2 mm; 40X scale bars: 25 µm). (**E**) Confocal images for assessment of sarcomere structure in animals six weeks following viral dosing (n = 4/condition, 90X scale bars: 25 µm). (**F**) Adult WT mice were dosed retro-orbitally IV with vehicle or the indicated doses (vg/kg) of AAV9:mMybpc3 and cardiac transgene expression was analyzed 10 weeks post-injection by qPCR for *Mybpc3* RNA (normalized to *Gapdh*). Protein assessment in WT mice was performed by ELISA with equivalent total protein for each sample, as well as (**G**) immunoblot (n = 5 – 6/condition). *p < 0.5, ** p <0.01, *** p <0.001, ****p <0.0001 per one-way ANOVA with Tukey’s multiple comparisons test. Data are shown as means ± SEM.

To determine the extent of cardiomyocyte transduction, immunohistochemistry for MYBPC3 four weeks post-injection was performed. Results indicated the ability to restore MYBPC3 protein expression to the majority of cardiomyocytes in *Mybpc3^−/−^* mice at both 3E13 and 1E14 vg/kg dose levels (Figure 5D). Thus, cardiac restoration of wild-type MYBPC3 protein levels corresponded to the comparable efficacy observed between *Mybpc3^−/−^* animals dosed with 3E13 and 1E14 vg/kg (Figure 4). Further, confocal imaged co-staining for α-actinin demonstrated proper sarcomeric incorporation in animals six weeks following viral dosing (Figure 5E).

To investigate the dose-dependent increase in transgene RNA without comparable increases in protein levels, we assessed transgene RNA and protein levels in a pilot safety evaluation of AAV9:mMybpc3 in naïve adult CD-1 mice. AAV9:mMybpc3 administered to WT mice was selected to achieve the greatest potential total levels of MYBPC3 protein expression. Retro-orbital IV injection at 10 weeks of age was used to deliver AAV9:mMybpc3 at 3E13 or 3E14 vg/kg, or vehicle control. AAV9:mMybpc3 was well tolerated in naïve CD-1 mice and cardiac function and morphology were unaffected by treatment and similar among all groups, and body weight remained stable (Figure S5).

Interestingly, AAV9:mMybpc3 administered at 3E14 vg/kg to WT animals exhibited a 6.5-fold increase in *Mybpc3* RNA (normalized to *Gapdh*) above vehicle-treated animals (Figure 5F), but no statistically significant increase in MYBPC3 protein (Figure 5F and 5G) based on ELISA. This observation of an increase in RNA without a proportional increase in protein is consistent with published findings that show transgenic mouse lines over-expressing *Mybpc3* RNA do not have MYBPC3 protein levels greater than WT^42,43^. Taken together, these results explain the comparable restoration of cardiac MYBPC3 protein for 3E13 and 1E14 vg/kg, despite significant differences in transgene RNA, and highlight the attractive safety profile of MYBPC3 gene replacement therapy, given apparent homeostatic regulation of protein expression to near WT levels *in vivo*.

### AAV9:mMybpc3 Improved Cardiac Diastolic Dysfunction at 3E13 vg/kg

Homozygous mice exhibited diastolic dysfunction manifested by slowed maximal mitral E-wave velocity (MV E), slowed mitral annular e’ velocity, and prolonged isovolumic relaxation times (IVRT) at 14 weeks of age (Figure 6 and Table S3). The decrease and prolongation in these parameters indicate heart muscle relaxation slowing. AAV9:mMybpc3 treatment was given systemically at two weeks of age via retro-orbital injections at 3E13 vg/kg and improvement of diastolic dysfunction was assessed 12 weeks later. As a control, vehicle was injected to a group of wild type (WT) and a group of *Mybpc3*^−/−^ animals (n=9/group). Importantly, AAV9:mMybpc3 was able to ameliorate diastolic dysfunction by improving these Doppler parameters in the *Mybpc3*^−/−^ mice. MV E treatment values were restored nearly to WT levels and were increased by 222 ± 44 mm/s relative to *Mybpc3*^−/−^ vehicle. The other diastolic parameters e’ and IVRT were improved by 7.2 ± 1.1 mm/s and 7.2 ± 1.2 ms, respectively, relative to vehicle. Taken together, the clinically relevant 3E13 vg/kg dose was able to ameliorate several diastolic dysfunction parameters associated with *Mybpc3*^−/−^ mice and HCM, and therefore partially restoring proper heart muscle relaxation.

**Figure 6.**
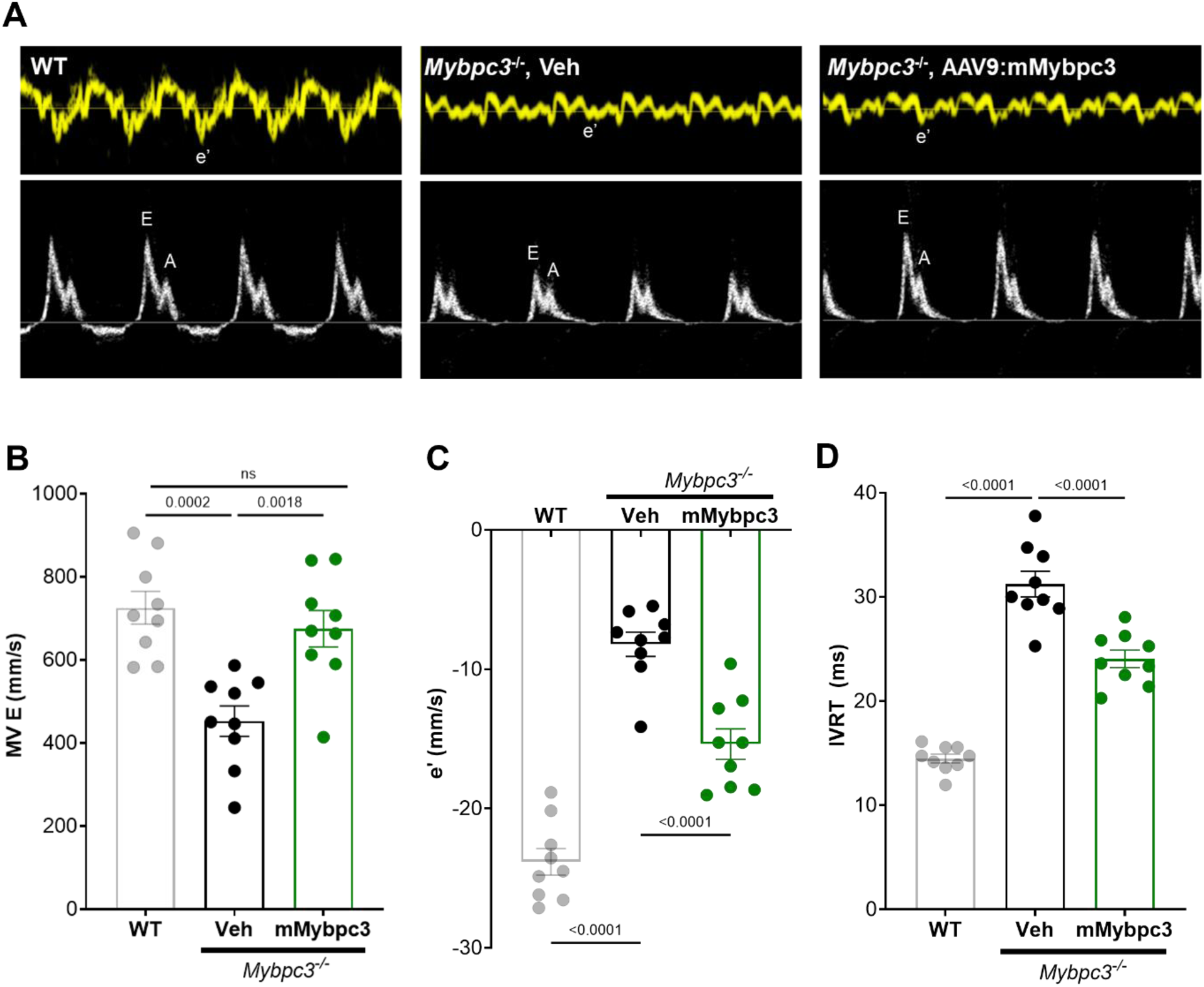
AAV9:mMybpc3 Improved Diastolic Dysfunction in *Mybpc3*^−/−^ Mice. Homozygous mice were dosed retro-orbitally IV with vehicle or 3E13 vg/kg of AAV9:mMybpc3 at two weeks of age and assessed for diastolic dysfunction 12 weeks post-injection. (**A**) Representative tissue (top) and pulsed-wave transmitral (bottom) Doppler tracings. Mitral annular e’ velocity, maximal mitral E-wave velocity (E), and maximal mitral A-wave velocity (A) shown. (**B**) Quantitation of mitral E-wave velocity (MV E), (**C**) mitral annular e’ velocity, and (**D**) isovolumic relaxation time (IVRT) demonstrated dysfunction in *Mybpc3*^−/−^ mice that was ameliorated by treatment, n = 9/group. P-value per one-way ANOVA with Tukey’s multiple comparisons test. Data are shown as means ± SEM

### Concomitant Decreased Markers of Fibrosis and Heart Failure

Having established significant improvements in cardiac function with a dose as low as 1E13 vg/kg, we sought to understand the molecular signature of MYBPC3 restoration in the dynamic dose range. Vehicle or AAV9:mMybpc3 at doses of 1E13 and 3E13 vg/kg were administered systemically via retro-orbital IV injection to symptomatic two-week-old *Mybpc3^−/−^*mice. An additional group of WT littermates were administered vehicle as a control. Echocardiography was performed to assess cardiac function until Week 14 post-delivery, at which point cardiac tissue was harvested for characterization.

Consistent with characterization of other *Mybpc3* mutant models,^44,45^ we observed significant upregulation of cardiac markers of heart failure (Figure 7A through 7C). Notably, the observed dose-dependency in echocardiography parameters (Fig. 4) was recapitulated in dose-dependent transcriptional reduction of cardiac markers of heart failure (Figure 7A through 7C), such as a BNP (*Nppb)*, where elevated levels are predictive clinically for HCM prognosis.^46^ Further, consistent with *MYBPC3*-associated HCM patients^47^ and other *Mybpc3* mutant models,^44,45,48^ we observed mild cardiac fibrosis which was beneath robust detection by trichrome staining (representative trichrome Figure S6 and whole heart images Figure S7 for 1E14 vg/kg study), and was clearly detected by cardiac transcriptional analysis as elevation of *Col3a1*, *Col4a1* and *Postn* transcripts (Figure 7D through 7F). As with heart failure markers, significant decreases in fibrotic gene expression were consistently observed in the AAV9:mMybpc3 3E13 vg/kg dosed animals compared to vehicle-treated *Mybpc3^−/−^* mice, whereas animals treated with 1E13 vg/kg exhibited only trending decreases.

**Figure 7.**
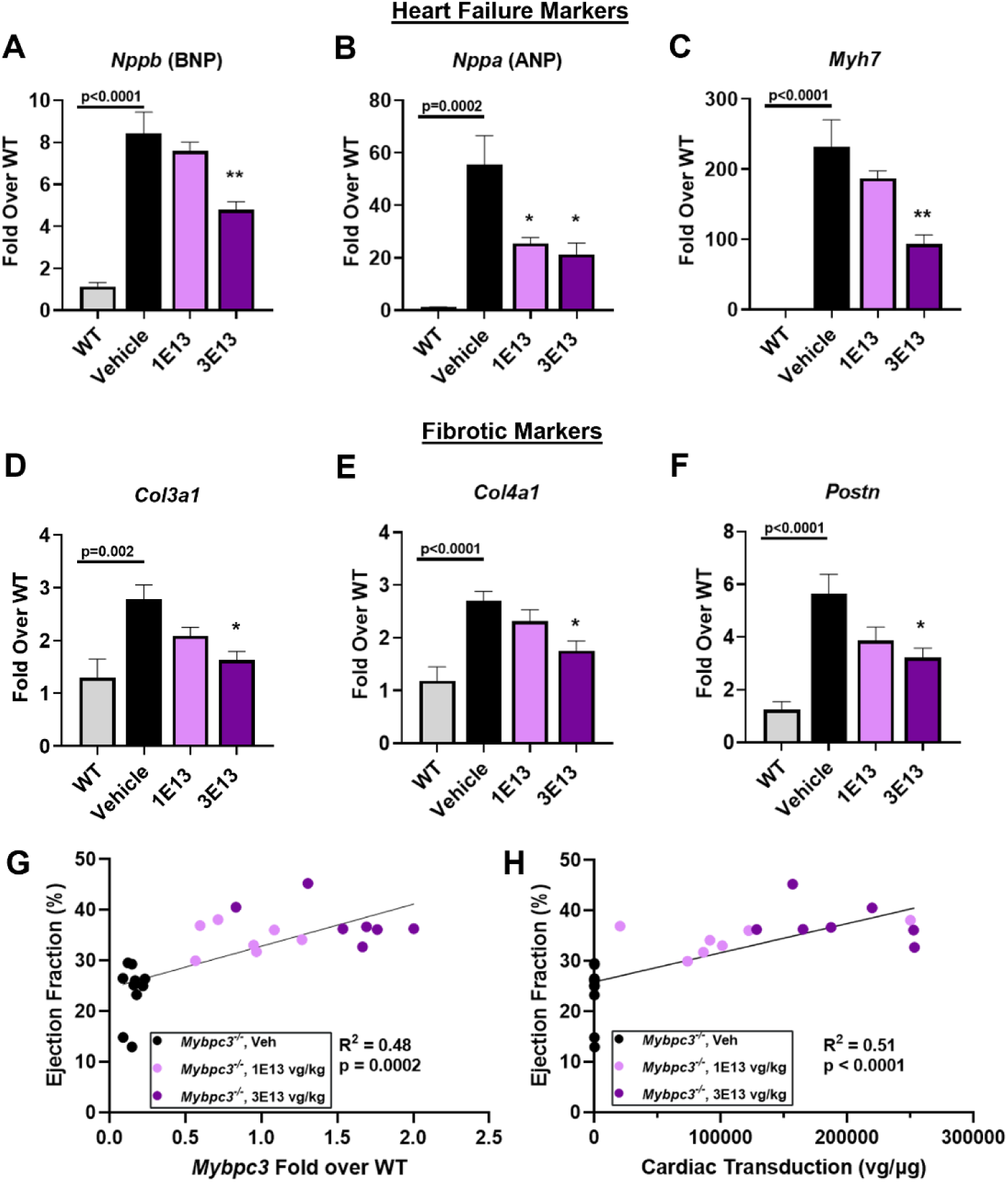
Dose-Dependent Inhibition of Expression of Genes Associated with Heart Failure and Fibrosis. Transcriptional analysis of cardiac tissue from homozygous mice dosed retro-orbitally IV with vehicle or the indicated doses (vg/kg) of AAV9:mMybpc3 at two weeks of age and WT littermates for heart failure markers (**A**) *Nppb*, (**B**) *Nppa*, and (**C**) *Myh7*. Dose-dependent decreases in fibrotic marker expression in treated animals as assessed for (**D**) *Col3a1*, (**E**) *Col4a1*, and (**F**) *Postn*. Significant correlation between ejection fraction and MYBPC3 restoration as analyzed by (**G**) transgene expression and (**H**) cardiac transduction. Analysis was performed 14 weeks post-injection (n = 7-10 mice/group). Data are shown as means ± SEM. * p < 0.5, ** p <0.01 compared to vehicle-treated animals per one-way ANOVA with Tukey’s multiple comparisons test.

To precisely determine how cardiac improvement mapped to restoration of MYBPC3 expression, we analyzed EF as a function of cardiac transduction and transgene *Mybpc3* expression in the individual *Mybpc3^−/−^* mice injected with AAV9:mMybpc3 at doses of 1E13 and 3E13 vg/kg. A dose-dependent increase in *Mybpc3* RNA expression was observed at the level of cardiac tissue from individual treated mice, and this transgene expression was significantly correlated with the extent of cardiac improvement observed (Figure 7G). Similarly, cardiac transduction exhibited dose-dependency which was positively correlated with EF (Figure 7H). Importantly, this demonstrated that consistent with analysis of the TN-201 cassette in human iPSC-CMs (Figure 1F through 1I), transduction of ≥ 1 vector genome per diploid genome (or ≥1.6E5 vg/µg DNA), as achieved with a dose of 3E13 vg/kg, resulted in MYBPC3 transgene expression equivalent to WT (Figure 7H and Figure 5A).

### Optimized Construct Outperforms Published Construct in Late-Stage *Mybpc3^−/−^* Mice

Rescue of function in symptomatic juvenile mice is, in the case of hypertrophic cardiomyopathy, more challenging than prevention of functional decline, as hypertrophic cardiomyopathy is a progressive disorder. As evidenced by the persistent slow decline and premature lethality of vehicle-treated *Mybpc3^−/−^*mice (Figure 3), older animals exhibit more severe disease than juveniles. Therefore, we sought to test the utility of cassette improvements in a more challenging model, homozygous adults with more advanced disease. To this end, we performed a side-by-side experiment comparing our optimized AAV cassette with a previously published 5.4 kb expression cassette encoding *Mybpc3* (Figure 8A), which had proven sufficient to prevent hypertrophy when injected into homozygous neonates.^36^

**Figure 8.**
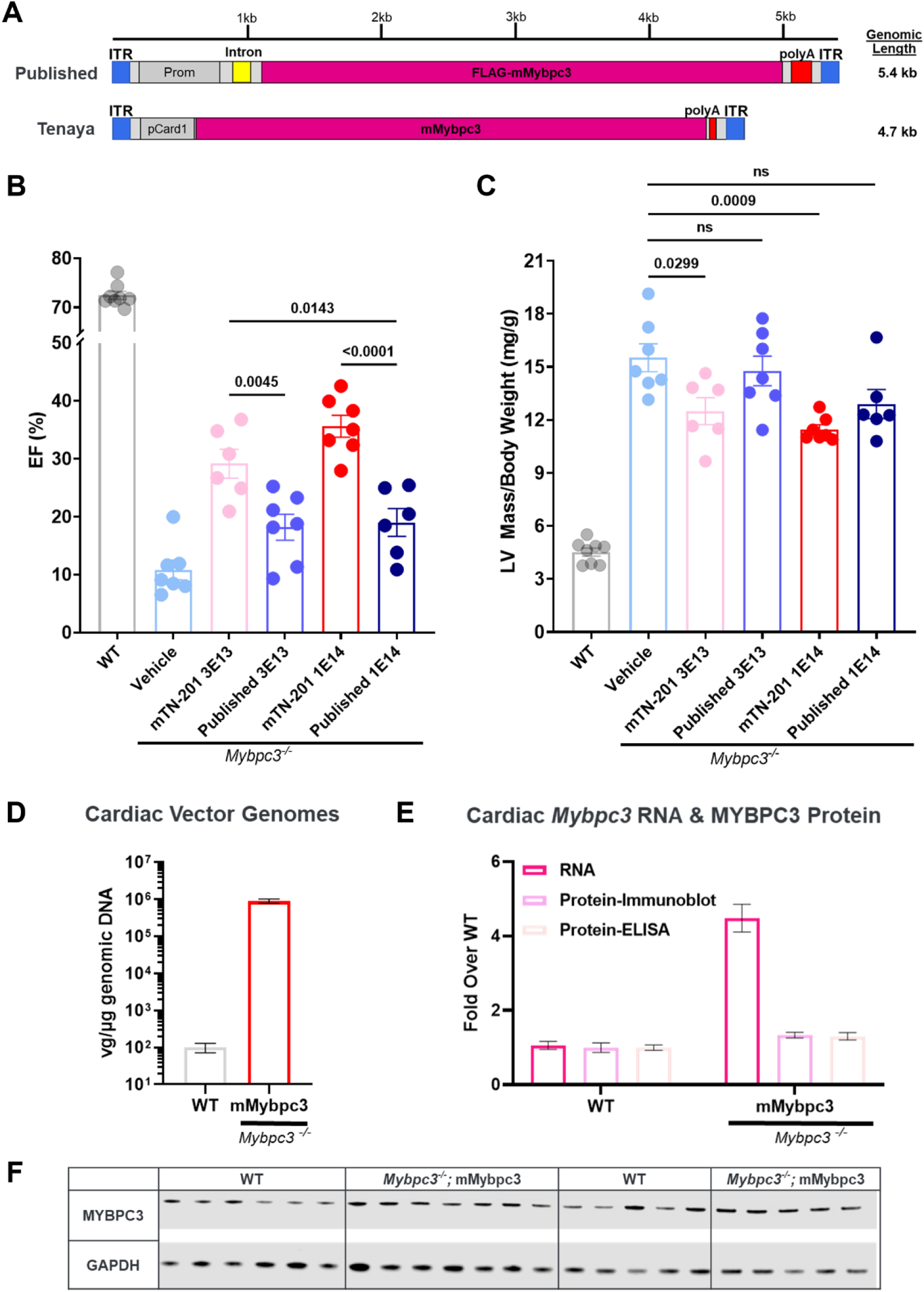
Optimized Construct Outperforms Published Construct in Late-Stage *Mybpc3^−/−^*Mice. (**A**) Homozygous mice with advanced disease (2.5 months of age) were injected retro-orbitally with 3E13 vg/kg or 1E14 vg/kg of AAV9 vector encoding *Mybpc3* in the context of the published 5.4 kb cassette (denoted “Published”) or the TN-201 mouse surrogate (AAV9:mMybpc3, denoted mTN-201), or injected with vehicle control. Both cassettes utilize sequence from the human cardiac human cardiac troponin T (*TNNT2*) promoter. (**B**) EF was measured to represent contractile ability at 27 weeks post-delivery. (**C**) LV mass was measured to represent hypertrophy and was normalized to body weight at 27 weeks post-delivery (n = 6-7 mice/group). (**D**) Sustained cardiac transduction 20 months post-injection drove (**E**) sustained cardiac *Mybpc3* RNA at ∼4X levels of endogenous transcript and sustained restoration of wild-type protein levels in dosed homozygotes 20 months post-injection. MYBPC3 protein expression was assessed by ELISA with equivalent total protein for each sample, as well as (**F**) immunoblot of equivalent total protein, with MYBPC3 intensities normalized to WT. P-value per one-way ANOVA with Tukey’s multiple comparisons test. Error bar: mean ± Standard Error of the Mean (SEM).

Homozygous mice with advanced disease (2.5 months of age) were injected retro-orbitally with 3E13 vg/kg or 1E14 vg/kg of AAV9 vector encoding *Mybpc3* in the context of the published 5.4 kb cassette (denoted “Published”) or the TN-201 mouse surrogate (AAV9:mMybpc3, denoted mTN-201) or injected with vehicle control. Strikingly, the combined cis-regulatory modifications of our cassette resulted in significantly greater improvements in EF than the published cassette at 3E13 vg/kg (29% compared to 18%) and at 1E14 vg/kg (36% compared to 19%). Further, the optimized cassette exhibited significantly greater EF improvement at a dose of 3E13 vg/kg than the published cassette at 1E14 vg/kg or a 3.3X increase in dose (Figure 8B). Improvements in cardiac hypertrophy exhibited a similar pattern, with the published cassette failing to have any significant effect on hypertrophy at the doses of 3E13 vg/kg and 1E14 vg/kg (Figure 8C). In contrast, the optimized cassette resulted in more significant improvements compared to vehicle-treated *Mybpc3^−/−^* mice at both doses tested (≥3.0 mg/g). Thus, the optimized cassette outperformed the published cassette for improvements in hypertrophy and cardiac function and resulted in statistically significant benefit even in this model of advanced *MYBPC3*-associated cardiomyopathy.

### Sustained Cardiac Transgene RNA and Protein Expression 20-months Post-Injection

Critical to the translational application of gene therapies is the durability of the AAV genome and therefore transgene expression in the cell type of interest, given current challenges that bar redosing. We therefore utilized the significant extension of lifespan afforded by AAV9:mMybpc3 in *Mybpc3^−/−^*mice to assess the persistence of cardiac transduction and transgene expression 20-months post-injection at a dose of 1E14 vg/kg. Consistent with durable efficacy findings for AAV9:mMybpc3 (Figure 3), terminal cardiac analyses indicated sustained cardiac transduction (Figure 8D), transgene RNA (Figure 8E) and restoration of wild-type levels of MYBPC3 protein 20 months post-injection. (Figure 8E and 8F).

## DISCUSSION

The efficacy of TN-201 has been demonstrated in a *Mybpc3^−/−^*mouse model, which develops marked LV hypertrophy, poor cardiac function, and dilation at two weeks of age, resulting in death around one year of age, similar to patients with HCM with bi-allelic truncating or null mutations. Demonstrating efficacy in this model is challenging due to the severe phenotype of the *Mybpc3^−/−^* mice and the lack of any MYBPC3 protein. In these studies, a single dose of TN-201 or AAV9:mMybpc3 (mouse surrogate of TN-201 incorporating the mouse *Mybpc3* gene instead of the human one) was systemically administered to *Mybpc3^−/−^* mice at doses ranging from 1E13 to 1E14 vector genomes per kilogram of body weight (vg/kg) at two weeks of age, after onset of cardiomyopathy. A single dose of AAV9:mMybpc3 in juveniles improved survival, cardiac systolic function, and hypertrophy in a dose-dependent manner, all of which were durable out to 13 months, the latest time point assessed by echocardiography. Diastolic dysfunction, an early hallmark of HCM^49,50^ was also ameliorated with AAV9:mMybpc3. Notably, sustained cardiac transgene RNA and MYBPC3 protein expression was observed 20 months post-injection of AAV9:mMybpc3 in *Mybpc3^−/−^* mice, the last time point measured.

Differences between AAV9:mMybpc3 and TN-201 treatments in *Mybpc3^−/−^* mice are consistent with the sequence divergence between the mouse and human orthologs (88% amino acid identity). Further, differences may be due to the fact that rodents use α-myosin heavy chain (αMHC, or fast twitch myosin), rather than βMHC (or slow twitch myosin) in mature ventricular cardiomyocytes.^51,52^ This may result in lower potency of the human MYBPC3 to restrict the activity of the more active fast twitch myosin,^14^ potentially dampening its ability to reduce hypercontractility and hypertrophy over time in the murine model. Despite this sequence divergence, TN-201 mediated significant efficacy in multiple parameters assessed in this mouse cardiomyopathy model providing proof of mechanism for gene replacement therapy.

Dose-ranging efficacy studies exhibited restoration of wild-type MYBPC3 protein levels and saturation of cardiac improvement at the clinically relevant dose of 3E13 vg/kg. In contrast, previous studies have been unable to restore wild-type MYBPC3 protein levels to homozygous mice,^36,53^ even when dosing AAV within days of birth. Numerous clinical evaluations indicate that the frequency of safety events increases with dose.^54–58^ Thus, the increased potency the Tenaya cassette was able to achieve is critical to the translational safety of the *MYBPC3* gene replacement approach, as comparable improvement in cardiac function with the published cassette would require a >3-fold increase in dosage. Importantly, we also demonstrated that the cardiac transduction-transgene expression relationship observed *in vivo* in mouse cardiac tissue was recapitulated in human iPSC-CMs.

Also key to the translatability of the gene replacement approach is the consideration of transgene overexpression and any resultant potential safety concerns. These experiments generate two key findings that mitigate this concern: 1) naïve mice dosed with 3E14 vg/kg, 30X an efficacious dose, exhibited no alterations in cardiac function or body weight, and 2) a 7-fold increase in RNA without a proportional increase in MYBPC3 protein. This suggests exquisite post-transcriptional regulation of MYBPC3 protein expression and is consistent with transgenic mouse lines over-expressing *Mybpc3* RNA exhibiting WT levels of MYBPC3 protein.^42,43^ This may be due to the maintenance of precise stoichiometry of proteins bound in the sarcomere, with degradation of unincorporated sarcomeric components, such as excess MYBPC3, as a homeostatic regulatory mechanism.

A potential limitation of this work is the increased severity of our *Mybpc3^−/−^* model relative to established models. Our model has worsened cardiac dysfunction and LV hypertrophy, which may in large part be due to the different genetic background (129/Sv vs C57BL/6). Work in other heart failure models have shown that strain background can play a key role in phenotype.^59,60^ There are also similarities between our model and established models, however. The MyBP-C3^t/t48^and cMyBP-C^−/-44^ mice both develop cardiac dysfunction and LV hypertrophy by juvenile age. Similar to ours, when aged, the cMyBP-C^−/−^ mice have a decline in cardiac function.^45^ The continued limitation of all these models has been their lack of true haploinsufficiency, which comprises the majority of *MYBPC3* HCM patients. Our model and others quickly develop hypertrophy, dilation, and cardiac dysfunction, indicative of a combined HCM and DCM phenotype that in humans only develops in compound heterozygous children or a minority of heterozygous “end-stage” HCM adult patients.

In contrast to the homozygotes, the *Mybpc3^+/−^* mice do not develop the expected human phenotype, given that they exhibit normal^44^ or only slightly decreased protein levels.^45,61,62^ Thus, the asymptomatic heterozygous mice are not a useful model for the 40% decrease and the resultant haploinsufficiency experienced in the human heterozygotes. We were able to restore WT expression to homozygous mutant mice at 3E13 vg/kg, providing evidence that TN-201 has the potential to restore the missing 40% of MYBPC3 protein expression in human heterozygous patients. Thus, the robust benefits in terms of survival, regression of hypertrophy and increases in EF seen in this severe disease model suggest that TN-201 may benefit patients across the full range of severity of *MYBPC3*-associated heart disease.

## Acknowledgments

We thank members of the Tenaya Animal Care Team: Xiaomei Song, Cristina Dee-Hoskins, Kevin Robinson, Yolanda Hatter, Jessie Madariaga, and Carolina Gomez.

## Author contributions

L.M.L., W.G.T., T.H., and K.I. conceived the idea. L.M.L., A.G.S, W.G.T, and T.H. designed the studies. L.M.L., T.N.Q, E.L., and A.G. generated and maintained the mice. A.G.S. performed all echocardiographic measurements and analyses, with assistance from J.Y.. S.S. and J.H. generated the iPSC line. A.B. and M.C. performed the iPSC-CM differentiations. C.F., T.W.C., N.G., S.J., Z.C., C.R., C.F. and E.E. generated the virus. C.A.L. and J.L. analyzed virus for release. F.J. and W.S.P. oversaw viral production and release. Z.C. and C.R. performed the viral injections. L.M.L., A.G., E.L., T.N.Q., K.K., and J.W. performed all in vitro experiments and ex vivo and analyses. L.M.L, A.G.S., E.L., T.N.Q., A.G., K.K., J.W., J.M.L., K.I., W.G.T and T.H. interpreted the results of the experiments. L.M.L wrote the manuscript with the review and support from all authors.

## Data and materials availability

All data associated with this study are included in the paper or the Supplementary Materials.

## SUPPLEMENTAL MATERIAL

### Animal studies

Animal studies were performed according to Tenaya Therapeutics’ animal use guidelines. The animal protocols were approved by the Institutional Animal Care and Use Committee (IACUC number: 2020.007).

### Mouse model

Applied Stemcell Inc. (Milpitas, CA) was contracted for creation of *Mybpc3* Exon1-Exon2 knockout mice, including generation of paired guide RNAs (GGCATCAAGCAGGCCACCCA and AAAGGTCAAGTTTGACCTCA) and utilization of the CRISPR/Cas9 system on a C57BL/6 background. Three male founders were obtained; TOPO cloning followed by sequencing was used to confirm the presence of the deletion in all founders. The germline transmission of the deletion was confirmed in F1 progeny by genotyping using forward primer (GAGAAGCCAGAGGACCAAGTG) and reverse primer (GGACCCTTCCTAGAACACCG, ∼376bp). The wild-type allele was assessed in parallel above forward primer and intronic reverse primer (GGAGCCAGGTCTCATGTGAA, ∼656bp). For genotyping, DNA was extracted from Viagen DirectPCR® Lysis Reagent (102-T) and proteinase K lysates by isopropanol precipitation. Three-primer PCR was run using Promega GoTaq® master mix. Breeders were maintained by backcrossing to C57BL/6 wild-type mice and experimental mice generated through timed heterozygous matings. Wild-type littermates were used as controls.

### Cardiac function

Cardiac function was assessed by transthoracic echocardiography using high resolution micro-imaging systems (Vevo 3100 Preclinical Imaging System, VisualSonics). Briefly, anesthetized spontaneously breathing mice (1-3% isoflurane and 98.5-99% O2) were placed in the supine position on a temperature-controlled heating platform to maintain their body temperature at ∼37°C. Nair was used to remove hair and expose the skin to the probe. Parasternal short-axis M-mode tracings of left ventricle (LV) were recorded for LV mass and LV ejection fraction (EF) calculations. EF was used to determine systolic function, whereas LV mass was used to determine hypertrophy. Vevo Lab software was used for analyses.

Simultaneous to echocardiography recordings, electrocardiography (ECG) leads were placed on animals in the Lead II position and ECG signals recorded with ADInstruments Powerlab. LabChart 8 software was used to assess QT intervals.

### Cardiac transgene and transduction analyses

RNA was extracted from cardiac tissue using the miRNeasy mini Kit (Qiagen Sciences, 217004), cDNA synthesized (Invitrogen SuperScript III First-Strand Synthesis SuperMix for qRT-PCR) and analyzed by qRT-PCR using the indicated Taqman probes listed in Table S4.

For cardiac lysates, equal masses of tissue were homogenized in RIPA Lysis Buffer and 1X Protease Inhibitor. A Pierce™ BCA Protein Assay Kit was run to determine the precise protein concentration of each sample lysate. Equal protein amounts were loaded for Western Blotting and detected with mouse anti-MYBPC3 (sc-137180, Santa Cruz Biotechnologies) and rabbit anti-GAPDH (ab181602, Abcam). Imaging was performed on the Li-Cor Odyssey and analyzed using ImageJ (NIH) densitometry analysis. Detection by ELISA was performed with MyBPC3 DuoSet ELISA (R&D Systems, DY7439-05).

For cardiac vector genomes analysis, DNA was extracted using the DNeasy Blood and Tissue Kit (Qiagen Sciences, 69506). Absolute quantification of cardiac viral genomes per microgram of genomic DNA was assessed by qPCR using linearized standards across six orders of magnitude. Primers used are listed in Table S5.

### Matched RNA, protein and transduction analysis in iPSC-CMs

As above, with cells being directly harvested from the well with Qiazol (Qiagen). DNA digestion was then performed using TURBO DNase (Invitrogen) and RNA was repurified using Quick-RNA MicroPrep kit (Zymo).

On-plate lysis was performed by harvesting iPSC-CMs 7-days post-infection with RIPA Lysis Buffer and 1X Protease Inhibitor, as above. Equal protein amounts were loaded for Western Blotting on a NuPage™ Tris-Acetate gel. Detection of MYBPC3 (sc-137180, Santa Cruz Biotechnologies) and Tubulin (ab7291, Abcam) was performed on the Li-Cor Odyssey.

For vector genomes analysis, cells were harvested directly from the well in Buffer ATL with proteinase K (Qiagen) then incubated at 56°C for 10 minutes at 1,000 RPM on a thermomixer. Lysates were then treated with 8 uL of RNase A (Qiagen) for 20 minutes at room temperature. DNA was extracted using equal volume of Phenol/Chloroform/Isoamyl alcohol (PCI) solution (25:24:1), with a chloroform back extraction prior to precipitation. DNA samples were quantified using the Qubit 1x dsDNA High Sensitivity Assay Kit (Life Technologies). To specifically quantify nuclear vector genomes, the DNA samples were normalized by concentration and treated with T5 exonuclease (New England Biolabs) at two different dilutions (1:5 and 1:50) to ensure enzyme was not limiting. T5 exonuclease is a single stranded DNA endonuclease and double-stranded DNA exonuclease, thus T5 digest can be used to determine the fraction of vector genome signal that is nuclear or episomal.^63,64^ For T5 treatment, 25 uL of each dilution was aliquoted for treatment and mock treatment. All samples were incubated at 37°C for two hours before being heat inactivated at 70°C for 10 minutes then analyzed by qPCR using 2x PowerUp™ SYBR™ Green Master Mix (Applied Biosystems) on a Quant Studio 7 (Applied Biosystems) with primers that specifically bind to the AAV genomic cassette, rather than transgene RNA, endogenous RNA or DNA. Quantification of nuclear viral genomes per microgram of genomic DNA was assessed by qPCR using linearized standards across six orders of magnitude. The lower limit of detection of the assay resulted in values of 1.92E3 as the background level of signal, equivalent to ∼1% of that detected in *MYBPC3^−/−^* iPSC-CMs dosed at 1K.

### Transient Transfection of iPSC-CMs

Confluent *MYBPC3^−/−^* iPSC-CMs were transiently transfected ten days after seeding using 500 ng of plasmid DNA in OPTI-MEM^TM^ (ThermoFisher) per 24-well. 2 µl of ViaFect^TM^ (Promega) in OPTI-MEM^TM^ was used per 24-well. ViaFect^TM^ and DNA solutions were then combined and complexes allowed to form for 20 minutes at room temperature. 50 µl transfection mixes were added dropwise to each 24-well and lysates harvested after seven days.

### Immunohistochemistry

The base of each heart was fixed overnight at 4C in 4% paraformaldehyde then dehydrated with progressive sucrose exchanges until sinking in 30%. Hearts were embedded in Optimal Cutting Temperature (OCT) compound and sectioned. Tissue was permeabilized and stained with the M.O.M.® (Mouse on Mouse) Immunodetection Kit (Vector Labs, BMK-2202), mouse anti-MYBPC3 (sc-137180), and rabbit anti-alpha Actinin 2 (PA5-27863).

### Flow cytometry

Cells were dissociated with TrypLE (Gibco), quenched, and assessed on a SH800 cell sorter (Sony Biotechnology).

### Statistical Guidelines

The number of technical and biological replicates and animals for each experiment are indicated in the figure legends. Statistical analyses were performed using Prism 9. P-values per one-way ANOVA with Tukey’s multiple comparisons test were utilized whenever more than two groups were being compared. Student’s t test was used to analyze two unpaired groups. Significant differences were defined as P < 0.05. Error bars in all in vivo studies represent SEM.

**Figure S1.**
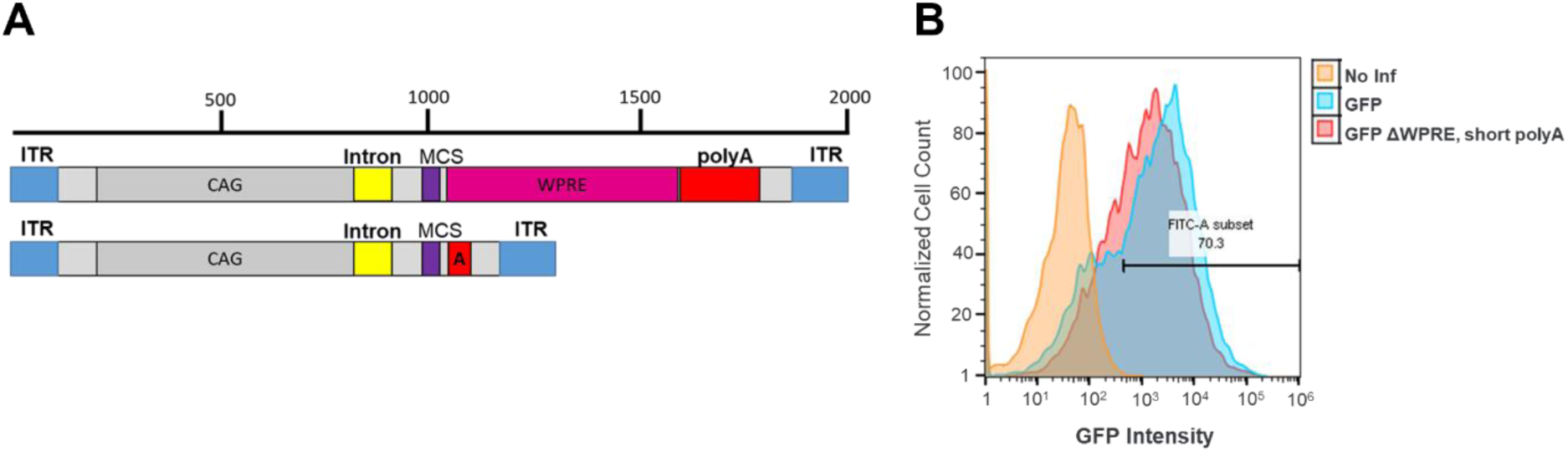
Minimized *cis*-regulatory AAV transgene cassettes. (**A**) Schematic of *cis*-regulatory cassette modifications. Starting with a standard AAV vector having a CAG promoter, intron, WPRE, and standard polyA sequence totaling ∼2 kb of regulatory sequence, we assessed the impact of woodchuck hepatitis virus posttranscriptional regulatory element (WPRE, 589 bp) removal, as well as polyadenylation signal abbreviation (“A”) to save an additional 170 bp. Multiple Cloning Site, MCS. (**B**) Expression of a GFP reporter cloned into the multiple cloning site (MCS), as assessed by flow cytometry analysis 2 days post-infection, indicated slightly decreased but robust expression in human cardiac fibroblasts (n = 2), MOI 160, 000.

**Figure S2.**
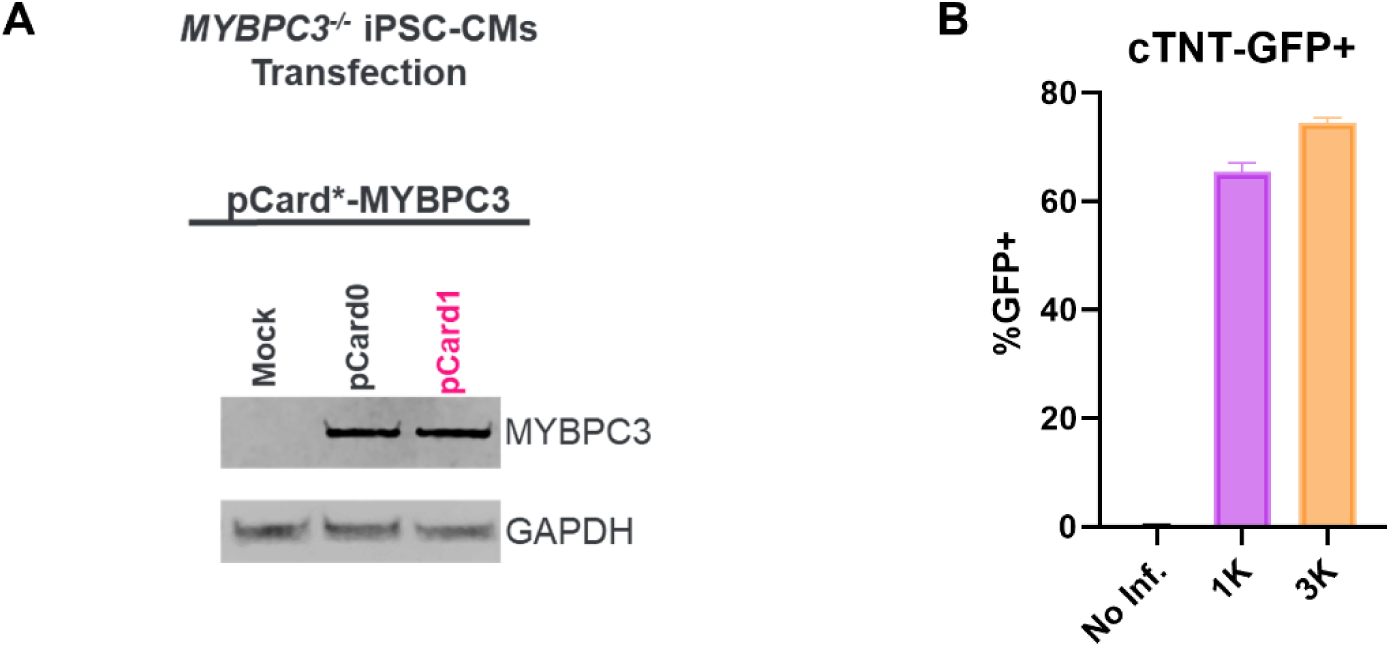
Promoter Assessment in Human iPSC-CMs. (**A**) The expression driven off of the unpackaged plasmid DNA of pCard0 and pCard1 constructs was assessed by transient transfection of *MYBPC3^−/−^* iPSC-CMs seven days after transfection. Consistent with the pCard1 4.7 kb version driving increased expression through improved packaging, no difference in protein expression was observable when *MYBPC3^−/−^* iPSC-CMs were transiently transfected with the naked plasmids. (**B**) Flow cytometry analysis of iPSC-CMs transduced with CR9-01:pCard1-GFP one week post-infection (N = 2/condition assayed). Consistent with vector genome analysis, flow cytometry one week post-infection with CR9-01:pCard1-GFP, a green fluorescent protein reporter, demonstrated that at the lowest dose used (1K) ∼60% of iPSC-CMs were transduced indicating an infectious multiplicity of infection equivalent to one. Data are shown as means ± SEM.

**Figure S3.**
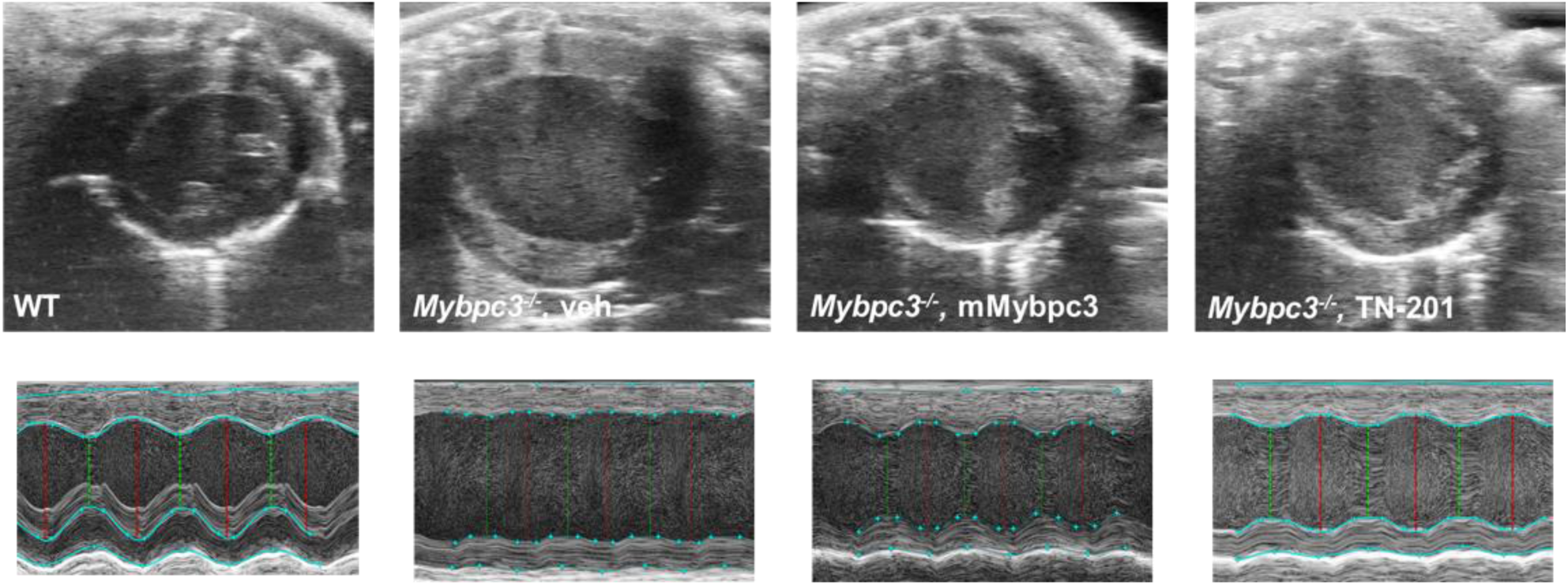
Representative B-mode (top) and M-mode (bottom) ultrasound images. WT (far left); *Mybpc3^−/−^* vehicle (center left); *Mybpc3^−/−^*, mMybpc3 (center right); and *Mybpc3^−/−^*, TN-201 (far right) shown.

**Figure S4.**
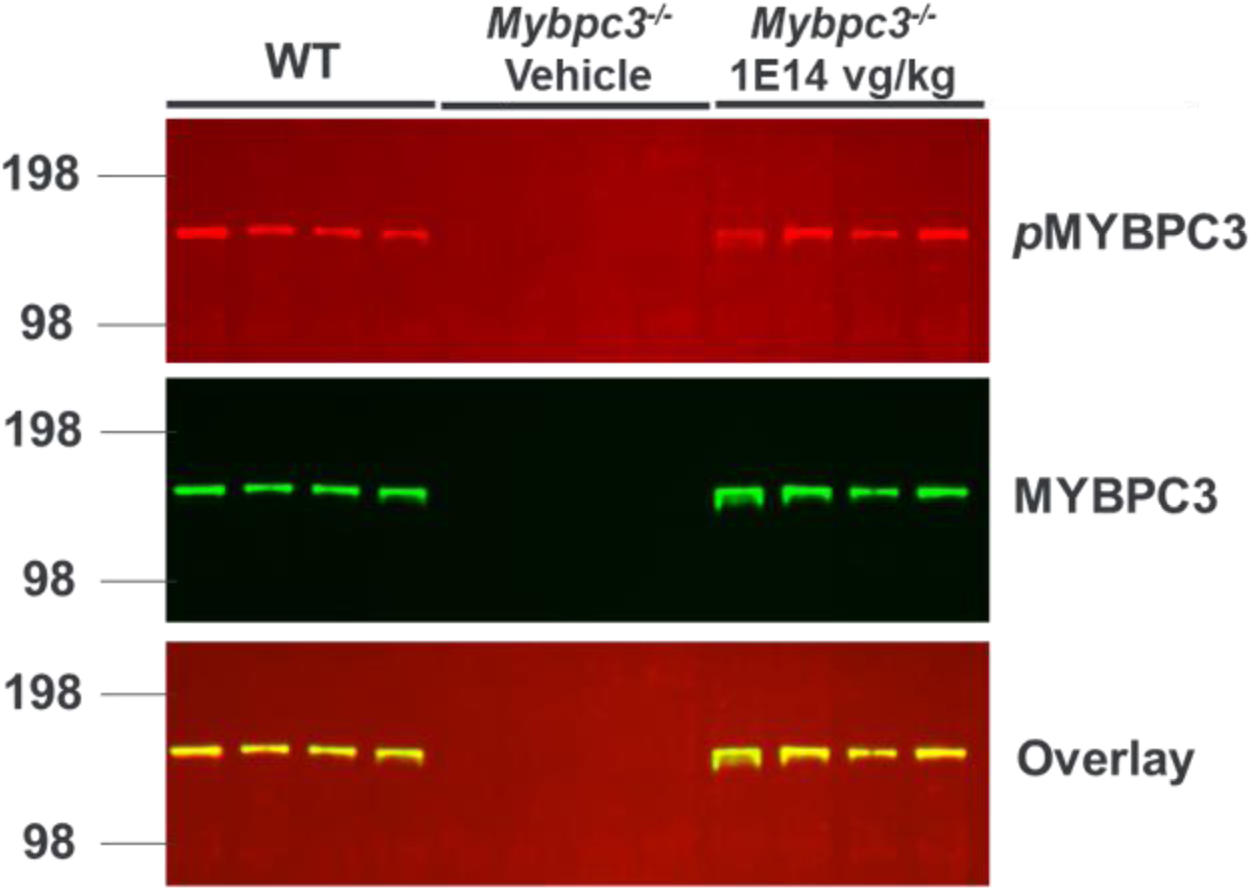
PKA-Mediated Phosphorylation of Restored Cardiac MYBPC3. Homozygous mice were dosed retro-orbitally IV at two weeks of age with vehicle or 1E14vg/kg of AAV9:mMybpc3 and cardiac protein analyzed six weeks post-injection. Age-matched WT littermates were used as controls. Following BCA quantification, equivalent levels of protein were loaded and analyzed by immunoblotting for Phospho-PKA Substrate (RRXS*/T*)^65^ (top) detected in the 680 channel and total MYBPC3 protein (middle) detected in the 800 channel. Overlay bottom, n = 4/condition.

**Figure S5.**
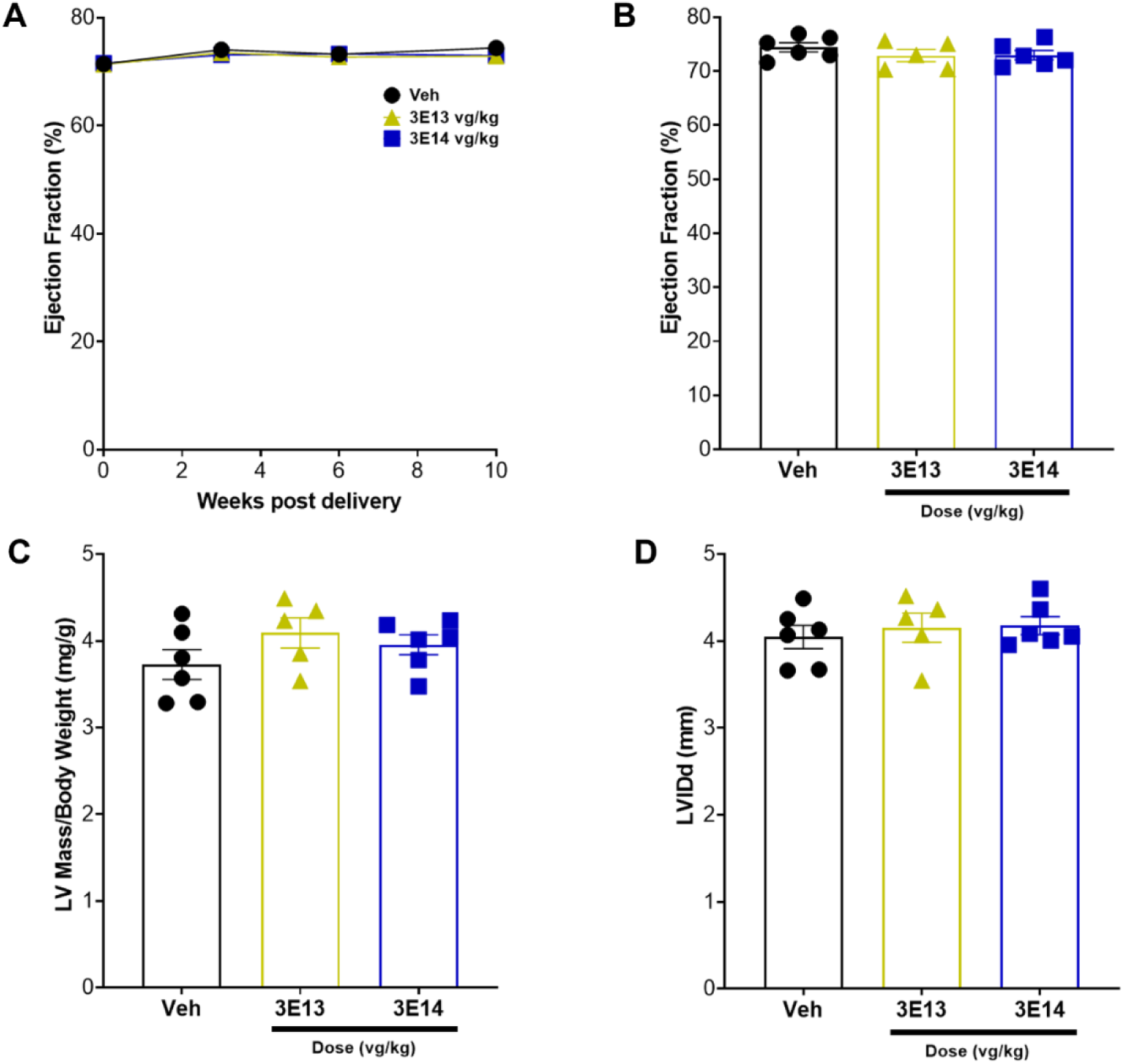
AAV9:mMybpc3 Did Not Affect Cardiac Function of Naïve Male CD-1 Mice at 30X an Efficacious Dose. (**A**) Ejection fraction (EF) progression and (**B**) EF at 10 weeks post-delivery. Gene delivery did not result in any changes in EF. (**C**) Left ventricular (LV) mass was measured to represent hypertrophy and was unchanged 10 weeks post-delivery. (**D**) Left ventricular internal diameter during diastolic (LVID;d) was measured to represent dilation and was unchanged 10 weeks post-delivery. Vehicle, Veh (n=6), 3E13 vg/kg (n=5) and 3E14 vg/kg (n=6).

**Figure S6.**
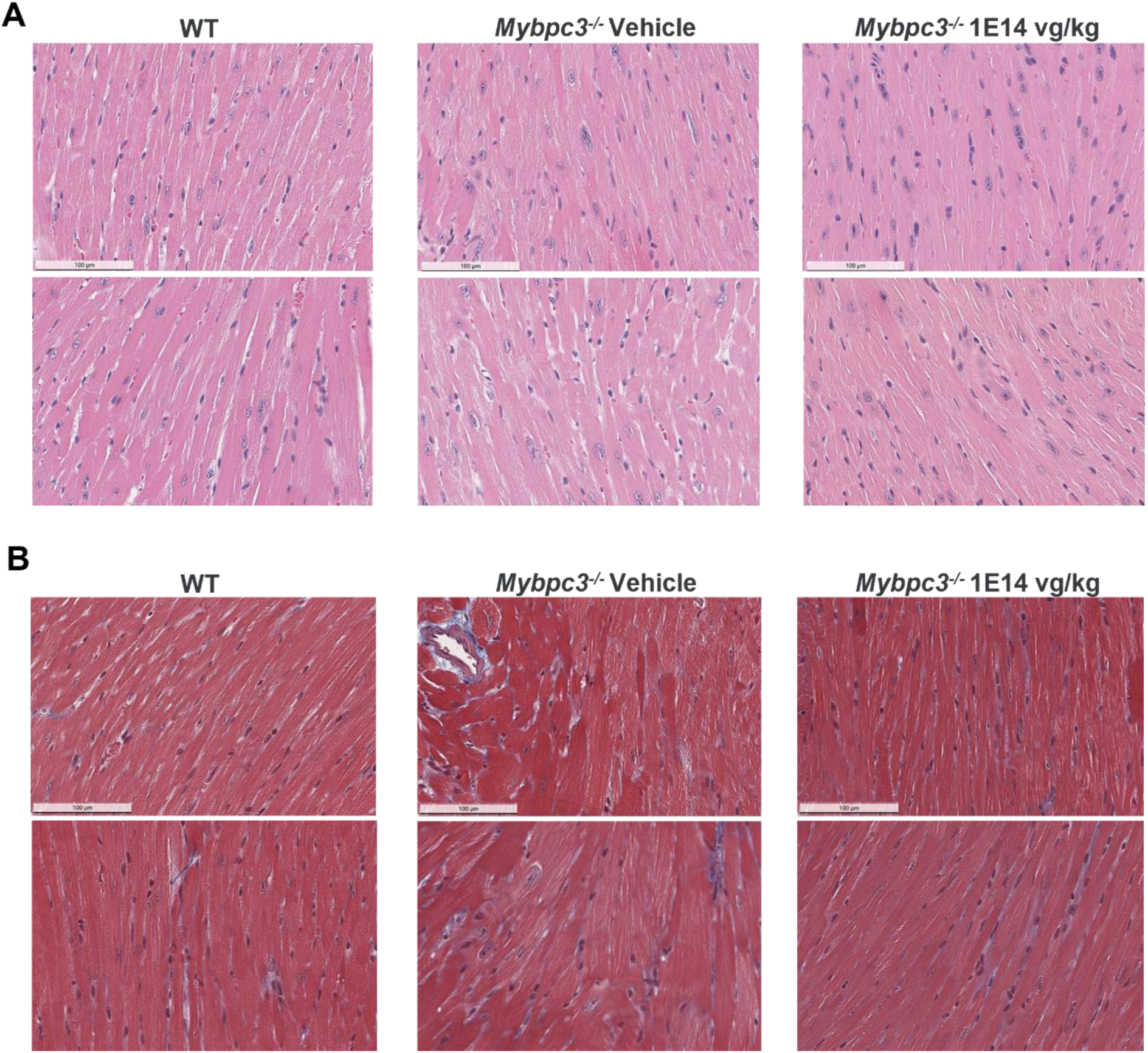
Representative H&E and Masson’s Trichrome Histology. Homozygous mice were dosed retro-orbitally IV at two weeks of age with vehicle or 1E14 vg/kg of AAV9:mMybpc3. Age-matched WT littermates were used as controls. Animals were euthanized at four months of age and their hearts fixed and paraffin embedded for (**A**) H&E staining and (**B**) Masson’s trichrome analysis.

**Figure S7.**
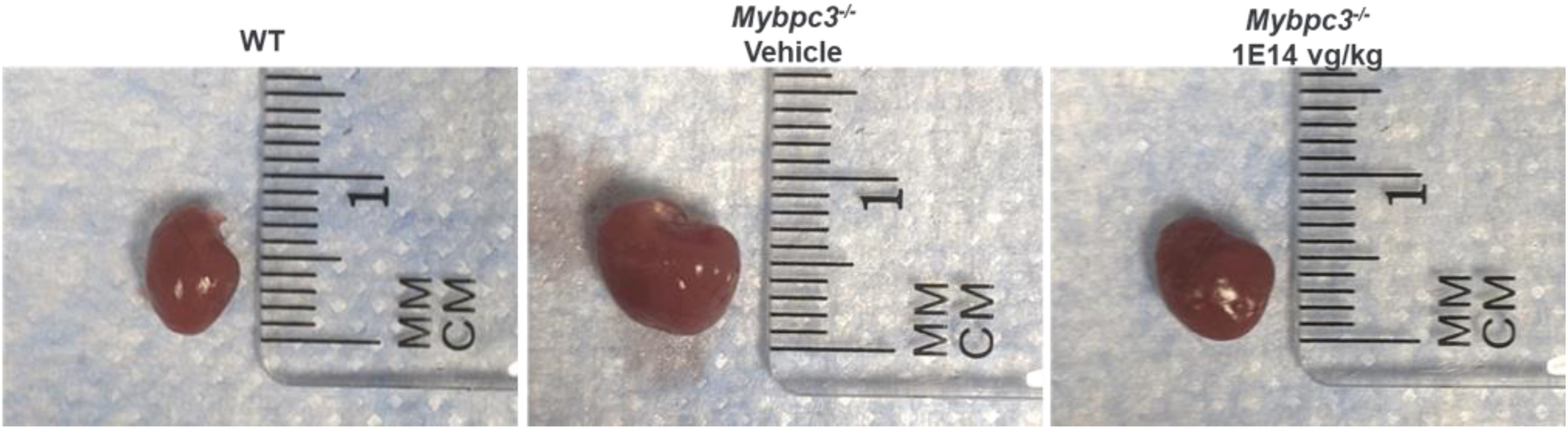
Representative Heart Images At Eight Weeks of Age. Homozygous mice were dosed retro-orbitally IV at two weeks of age with vehicle or 1E14 vg/kg of AAV9:mMybpc3. Age-matched WT littermates were used as controls. Animals were euthanized at eight weeks of age, or six weeks following viral injection.

**Table S1.** Echocardiography Parameters for Figure 2.

|  | WT | *Mybpc3*<sup>+/-</sup> | *Mybpc3*<sup>-/-</sup> |
| --- | --- | --- | --- |
| HR (bpm) | 509.0±19.8 | 496.3±18.4 | 493.0±9.0 |
| LVID;s (mm) | 1.57±0.04 | 1.63±0.03 | 3.22±0.08**** |
| LVID;d (mm) | 2.64±0.05 | 2.80±0.15 | 3.85±0.08**** |
| SV (μL) | 18.9±0.9 | 22.1±1.0 | 22.5±1.0* |
| EF (%) | 73.4±0.9 | 74.4±0.4 | 35.2±1.0**** |
| FS (%) | 40.7±0.7 | 41.7±0.3 | 16.5±0.5**** |
| LV Mass (mg) | 29.0±2.0 | 35.6±1.4 | 110.2±6.9**** |
| LV Mass/BW (mg/g) | 3.4±0.3 | 4.0±0.2 | 12.8±0.8**** |
| LVPW;d (mm) | 0.54±0.03 | 0.64±0.03 | 0.85±0.04** |

Heart rate (HR), left ventricular inner diameter during systole (LVID;s), left ventricular inner diameter during diastole (LVID;d); stroke volume (SV), ejection fraction (EF), fractional shortening (FS); body weight (BW), left ventricular poster wall thickness during diastole (LVPW;d). \*p<0.05, \*\*p<0.01, \*\*\*p<0.001, \*\*\*\*p<0.0001 relative to WT. Data shown as Mean ± SEM.

**Table S2.**
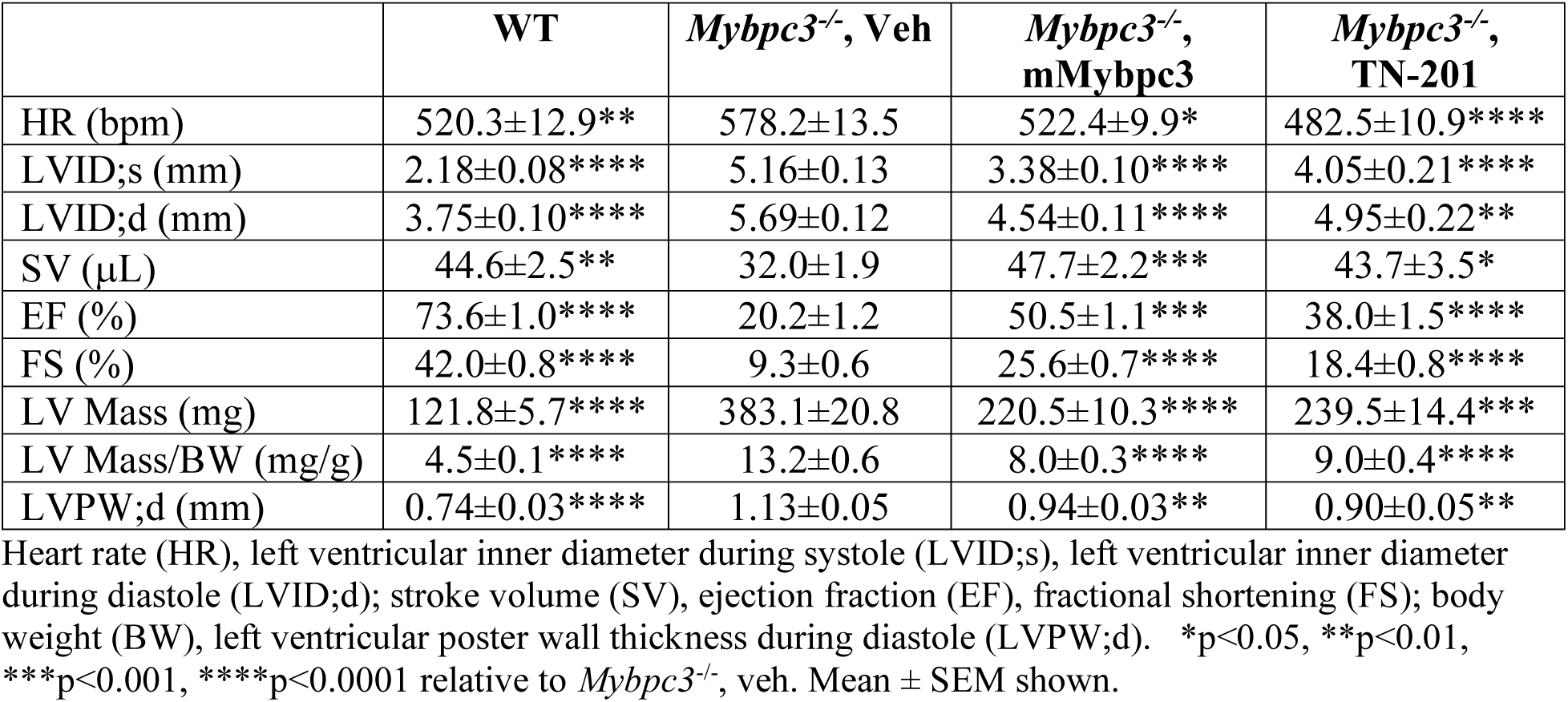
Echocardiography Parameters for Figure 3.

**Table S3.** Echocardiography Parameters for Figure 6.

|  | WT | *Mybpc3*<sup>-/-</sup>, Veh | *Mybpc3*<sup>-/-</sup>, m*Mybpc3* |
| --- | --- | --- | --- |
| e' (mm/s) | -23.8±1.0**** | -8.2±0.9 | -15.4±1.1**** |
| MV E (mm/s) | 725.6±39.3*** | 452.7±36.8 | 675.2±44.0** |
| MV A (mm/s) | 385.2±38.8 | 413.3±43.2 | 347.2±51.5 |
| IVRT (ms) | 14.5±0.4**** | 31.2±1.2 | 24.1±0.8**** |
| MV E/A | 2.02±0.20 | 1.14±0.06 | 2.55±0.60* |
| MV E/E' | -31.1±2.6*** | -57.9±5.6 | -45.2±3.7 |

Mitral Annular e' Velocity (e'), Maximal Transmitral E-wave velocity (MV E), Maximal A-wave velocity (MV A), Isovolumic Relaxation Time (IVRT) \*p<0.05, \*\*p<0.01, \*\*\*p<0.001, \*\*\*\*p<0.0001 relative to *Mybpc3*<sup>-/-</sup> veh. Mean ± SEM shown.

**Table S4.**
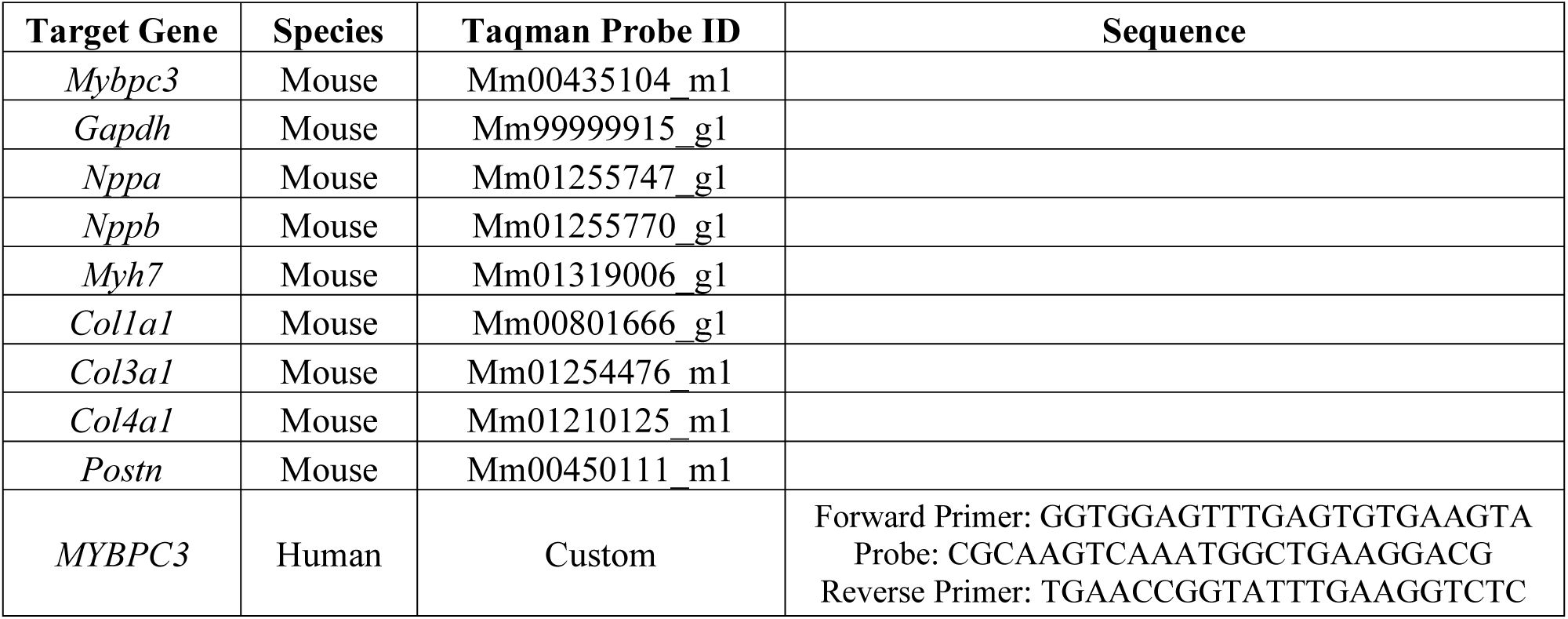
Taqman Probes for RNA Detection.

**Table S5.**
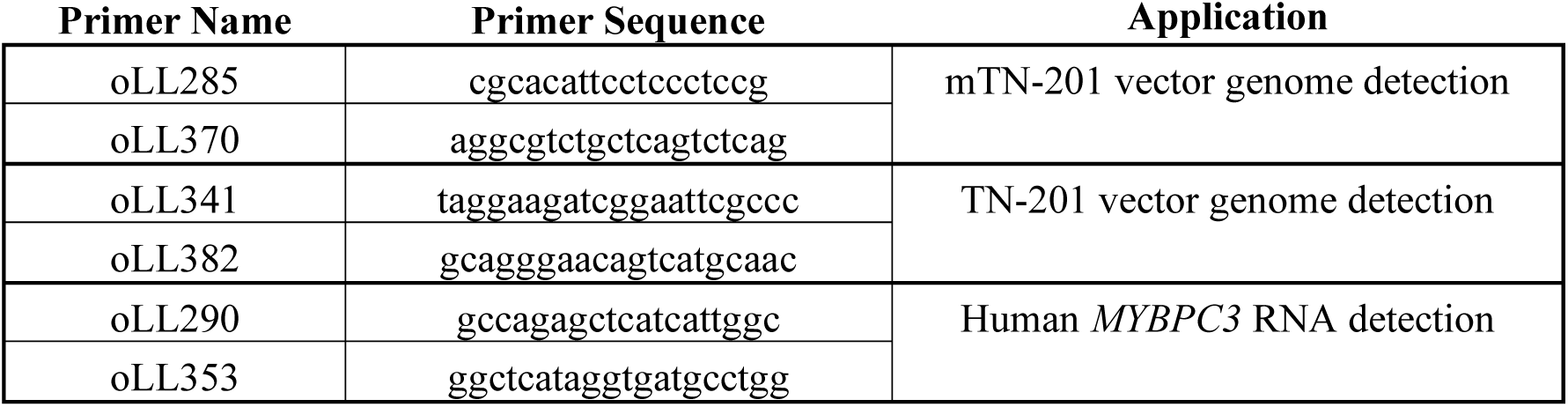
Primers Used With SYBR Detection.

**Table S6.** Viral Lots.

| **Virus** | **AKA** | **Lot** | **Figure** |
| --- | --- | --- | --- |
| AAV6: pCard0-MYBPC3 |  | CR #107 | 1C |
| AAV6: pCard1-MYBPC3 |  | 014VC19AUG28-1 | 1C |
| AAV6: pCard2-MYBPC3 |  | 014VC19AUG28-2 | 1C |
| CR9-01: pCard0-MYBPC3 |  | CR012820-142 | 1D |
| CR9-01: pCard1-MYBPC3 |  | CR012820-143 | 1D |
| AAV9: pCard0-MYBPC3 |  | 012VC19Aug07-1 | 1E |
| AAV9: pCard1-MYBPC3 |  | 021VC19Oct02 | 1E |
| CR9-01: pCard1-MYBPC3 | TN-201* | AP042822 | 1F-1I |
| CR9-01: pCard1-GFP |  | HZ011121 | 1F-1I |
| CR9-01: pCard1-MYBPC3 | TN-201* | EE061022 | 1A |
| TN-201 |  | 002VC08Jan2020-3 | 3 |
| AAV9:mMybpc3 | mTN-201 | 002VC08Jan2020-5 &<br>002VC08Jan2020-7 | 3, 5D, 8D-8F |
| AAV9:mMybpc3 | mTN-201 | 006VC19Feb2020-4 &<br>007VC26Feb2020-1 | 4, 5A-5C, 5E-5G,<br>S4, S5, S7 |
| AAV9:mMybpc3 | mTN-201 | ZC #821 | 6 |
| AAV9:mMybpc3 | mTN-201 | 009VC24Apr2020-1 | 7 |
| AAV9:mMybpc3 | mTN-201 | 020VC02Sep2020-9 | 8B, 8C |
| AAV9:mMybpc3 5.4kb | Published | 020VC02Sep2020-6 &<br>021VC09Sep2020-7 | 8B, 8C |
| AAV9:mMybpc3 | mTN-201 | PD210608 | S6 |

**Table S7.** Supplemental Immunoblot Quantification.

| **Blot** | **Construct** | **MYBPC3/GAPDH ± SEM** |
| --- | --- | --- |
| 1C | pCard0 | 0.54 |
| 1C | **pCard1** | **0.95 ± 0.37** |
| 1C | pCard2 | 0.40 ± 0.02 |
| 1D | pCard0 | 0.59 ± 0.00 |
| 1D | **pCard1** | **1.09 ± 0.02** |

